# Convergent and selective representations of pain, appetitive processes, aversive processes, and cognitive control in the insula

**DOI:** 10.1101/2025.02.18.638889

**Authors:** Mijin Kwon, Ke Bo, Rotem Botvinik-Nezer, Philip A. Kragel, Lukas Van Oudenhove, Tor D. Wager, The Affective Neuroimaging Consortium

**Author notes:** Corresponding authors: Tor D. Wager, Diana L. Taylor Distinguished Professor Presidential Cluster in Neuroscience and Department of Psychological and Brain Sciences Dartmouth College, Mijin Kwon, Department of Psychological and Brain Sciences Dartmouth College. Group author (The Affective Neuroimaging Consortium).

## Abstract

Regions that respond to multiple types of information (“convergence zones”) are crucial for the brain to generate coherent experiences and behaviors. The insular cortex, known for its functional diversity, has been hypothesized to be a key convergence hub, yet empirical evidence identifying how and where convergence occurs is incomplete. To address this gap, we analyzed functional convergence across four key task domains—somatic pain, non-somatic appetitive processes, aversive processes, and cognitive control—in a large-scale Bayesian mega-analysis of fMRI data (N=540, systematically sampled from 36 studies). Bayes Factor analyses identified both convergent zones responding to multiple domains and selective zones responding to single domains. Results revealed a hierarchical convergence architecture, with a multi-domain convergence zone in bilateral dorsal anterior insula surrounded by zones showing progressively increasing convergence. Functional decoding and coactivation analyses further support the insula’s role as a convergence hub, while cytoarchitectonic and neurotransmitter profiling characterizes the potential neuroanatomical underpinnings subserving convergent and domain-selective function. Overall, our results demonstrate a structured functional topography in the insula that bridges specialized and convergent processing, providing a potential neural basis for how diverse information streams combine into unified subjective experiences.

## Introduction

The human brain possesses an extraordinary ability to integrate a wide array of information from the body and environment into a seamless, unified subjective experience. The insula is thought to be central to this capacity, operating as a convergence zone—a region where diverse information streams are integrated—for a remarkably diverse range of interoceptive and exteroceptive processes, including visceral, autonomic, and homeostatic signals^1,2^, along with information from somatosensory, olfactory, gustatory, and auditory sensory inputs^3–7^.

However, most empirical research has focused on identifying subregions specific to particular aspects of sensation, emotion, and cognition^1–4,8–23^, documented across analyses of human neuroimaging^1–4,8–23^, pathways in non-human animals^24–27^, and human brain stimulation and electrophysiology^28–33^. In contrast, the concept of functional convergence zones in the insula has been largely theoretical^34,35^.

Convergence zones are thought to exist at multiple levels throughout the brain, from modality-specific integration to higher-order multi-modal convergence^36,37^. Of particular interest are zones that integrate external sensory information with internal states, as this integration is thought to be fundamental for constructing the sense of self^34,38^. The insula has been proposed as one potential site for such integration, with an influential theory by Bud Craig^38,39^, positing that the anterior insula (AIns) constitutes a convergence zone contributing to subjective, conscious experience. This aligns with concepts of embodied cognition, where the integration of interoceptive states and sensorimotor capacities underlies cognitive and affective processes and vice versa^40,41^.

Several lines of evidence support this view. The insula is one of the most functionally diverse regions of the brain^42^, integrating information at long time scales (several seconds or longer^43,44^), and coordinating functional relationships across brain networks^45^. Also, different insular neuronal populations encode diverse interoceptive and special sensory inputs, including visceroception^46,47^, immune afferents^25,48–50^, heartbeat perception^51–53^, pain^54–57^, and taste and smell^58,59^. Von Economo Neurons (VENs) located in AIns might provide a cellular substrate for rapid information integration^60,61^.

Despite the prominence of Craig’s theory, only a few studies have directly evaluated the concept of multi-modal convergence zones in AIns, and the precise locations of such zones have not been firmly established. To date, available research has relied on evidence of functional co-localization derived from Coordinate-Based Meta-Analyses (CBMAs) of the functional neuroimaging literature^4,19,20^, including Kurth et al.^4^, which identified partial overlap in AIns across functional domains. However, while informative, CBMA suffers from coarse spatial resolution as it uses activation coordinates representing only a fraction of activated voxels extracted from published results and applies smoothing to these peak locations, which are subject to considerable spatial noise^62^. This approach can produce artificial evidence of functional convergence where none exists and cannot systematically evaluate functional specificity across domains or test effect sizes across the entire insula in an unbiased manner.

To provide a comprehensive test of functional convergence in the insula and identify both convergent and selective functional zones, we conducted a mega-analysis of participant-level fMRI contrast maps selectively sampled from the Affective Neuroimaging Consortium (www.anic.science) database. This analysis focused on four functional domains closely linked to insular function: somatic pain, non-somatic appetitive processes, non-somatic aversive processes, and cognitive control^63^. We systematically included three subdomains per domain (e.g., three distinct types of somatic pain), with three studies per subdomain and 15 participants per study (Supplementary Fig. 1; *k*=36 studies, *n*=540). This design enabled us to test whether insular subregions encode domain information in a generalizable way across studies, which is critical for construct validation^64^. For example, general pain-encoding regions should consistently respond to different types of pain (e.g., thermal, mechanical, and visceral pain), while not responding to non-nociceptive perturbations (e.g., emotionally arousing stimuli). With these aggregated data and voxel-wise calculation of Bayes Factors^65^, we could directly test for both presence and absence of domain-specific activation at the voxel level, overcoming the spatial precision limitations of CBMA. This approach enabled systematic identification of insular subregions that: (1) activate across all domains, potentially indicating multi-domain convergence zone (“domain-general”), (2) activate selectively to specific functional domains (“domain-selective”), and (3) show gradients of cross-domain convergence.

Our analysis identified both multi-domain convergence zones in bilateral dorsal anterior insula (dAIns) and domain-selective zones for each domain, with a spatial organization where domain-selective inputs, processed separately in distinct insular subregions, progressively converge toward multi-domain zones in bilateral dAIns. However, shared responses across domains could reflect either true functional convergence or merely parallel but independent processes occurring in the same region. To distinguish between these possibilities and better understand the nature of these zones, we further characterized them through multiple levels of analysis: meta-analytic functional decoding (using Neurosynth^19,42^), coactivation with other brain regions^66,67^, cytoarchitectonic mapping^23,68^, and neurotransmitter system profiling^69,70^. Together, these findings provide support for both considerable functional diversity and multi-modal convergence zones in specific parts of the insula, especially in dAIns.

## Results

### Multivariate classification reveals inter-study generalizability across four functional domains

First, we tested whether each functional domain has a separable insular representation consistent across subdomains and studies, despite the heterogeneity introduced by aggregating data across different studies. We trained multiclass linear Support Vector Machine (SVM) classifiers to distinguish each domain from the others based on the multivariate activation patterns in the insula, holding out entire studies across 100 train-test splits stratified by subdomain to ensure generalizability across different experimental contexts. We report performance results averaged across these stratified samples (see Methods for details).

Domain-level classifier achieved above-chance prediction accuracies for three of the four domains (chance=25%): pain (74.71%; range: 40–100%; *p<0.001*), appetitive processes (60.73%; range: 40–80%; *p<0.001*), and cognitive control (52.00%; range: 17.78–75.56%; *p<.001*). Aversive processes showed lower accuracy (33.17%; range: 4.44–60%; *p=0.1327*). Examining the confusion matrices (Fig. 1b) indicated that aversive processes were confusable with appetitive processes (but less confusable with pain and cognitive control; see Supplementary Fig. 2 for pairwise predictive accuracy and further interpretation of the results), whereas pain was most discriminable from all other domains. Additional subdomain-level classification analyses revealed that some subdomains showed particularly distinct neural patterns (Supplementary Fig. 3), especially responses to aversive sounds (a subtype of aversive processes) and working memory (a subtype of cognitive control; see Study design in Methods for further description of subdomains).

**Fig 1.**
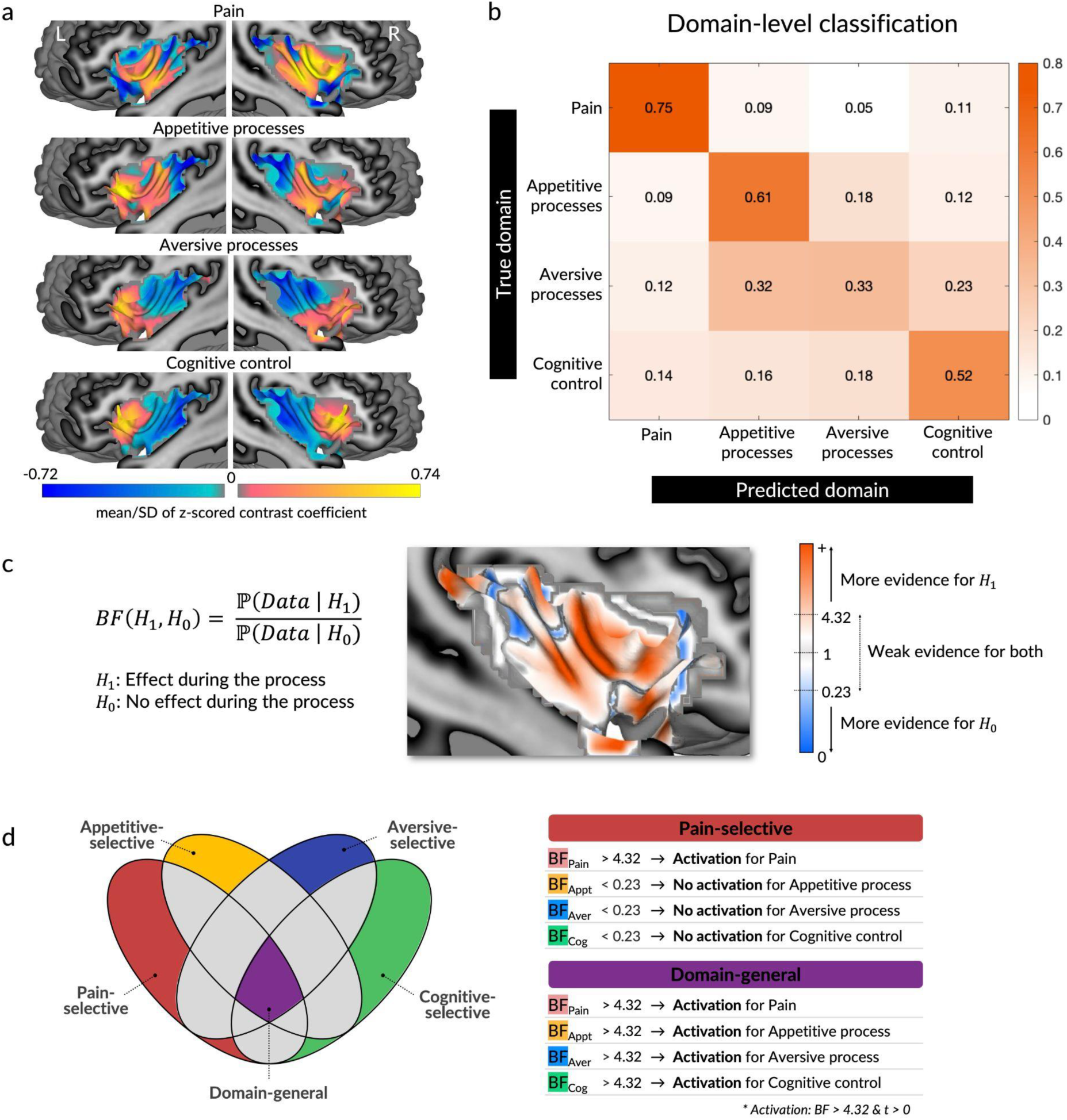
Multi-class SVM classification across four functional domains. **a,** Standardized insular activation pattern for each domain. Each contrast image was z-scored to ensure voxel weights to be on the same scale to facilitate cross-study harmonization and used for training SVM classifiers and subsequent analyses. Plotted values show the standardized means (mean contrast values across participants divided by the standard deviation. **b,** Prediction accuracies for SVM classifiers trained to discriminate between domains using a leave-one-study-out scheme. Above-chance prediction accuracies (25%) indicate separable and reliable neural representations for the given domain within the insula, despite the heterogeneity introduced by aggregating data across different studies. **c,** Identifying domain-general and domain-selective voxels using Bayes Factors (BFs). BFs were calculated for each domain to compare the probabilities of observing an effect versus no effect. BFs>1 indicate more evidence for an effect, while BFs<1 indicate more evidence for a null effect. Thresholds for sufficient evidence were set at 4.32 for effect (equivalent to FDR-corrected q<0.01) and 0.23 (the inverse of 4.32) for no effect. **d,** Definition of domain-general and domain-selective voxels. Voxels were considered activated only when BF>4.32 and t-statistics were positive and cases of either more evidence for no effect (BF<4.32) or evidence for deactivation (BF>4.32 with negative t-statistics) were considered no activation. Example map shows voxel-wise BFs for pain (red: stronger evidence for effect; blue: stronger evidence for no effect). Domain-general voxels were defined as those showing activation in all domains. Domain-selective voxels showed activation in their designated domain and evidence favoring no activation in other domains.

Overall, these findings indicate that three of the four domains show coherent insular representations (i.e., their activity patterns are consistent and distinguishable to enable significant above-chance classification across subdomains and studies). This coherence across different experimental paradigms and study conditions suggests reliable domain-specific neural representations. These results provide a foundation for identifying domain-general and domain-selective representations in individual regions.

### Domain-general and domain-selective regions identified using Bayes Factors

To identify insular voxels encoding domain-general or domain-selective responses, we used Bayes Factors (BFs) to assess both presence and absence of domain-specific activation *using* a Bayes Factors one-sample t-test^65^ (*n=135* per domain). We set a threshold of 4.32:1 odds favoring an effect versus no effect, corresponding to *q<0.01* FDR correction on average across domains (Fig. 1c; see Methods for details). Voxels were considered activated only when showing both *BF>4.32* and positive t-statistics. Evidence for no activation included cases of either more evidence for no effect (*BF<4.32*) or evidence for deactivation (*BF>4.32* with negative t-statistics). Based on these criteria, voxels activated in all four domains independently were classified as domain-general (Fig. 1d: purple), and voxels activated in one domain and showing evidence favoring no activation in the other domains were classified as single domain-selective (Fig. 1d: pain in red, appetitive in yellow, aversive in blue, and cognitive in green). For brevity, we refer to zones selective for each domain as pain-selective, appetitive-selective, aversive-selective, and cognitive-selective throughout the paper.

As shown in Fig. 2a, we identified specific regions within dAIns bilaterally that were activated across all domains and were therefore classified as domain-general. These zones included 120 of 6012 insular voxels and spanned anterior short and inferior gyri^71^. This part of dAIns maps onto regions most frequently activated across different task domains in the Neurosynth database^42^ and is substantially higher in the principal cortical gradient identified by Margulies et al.^72^, indicating transmodal (as opposed to unimodal) function.

**Fig 2.**
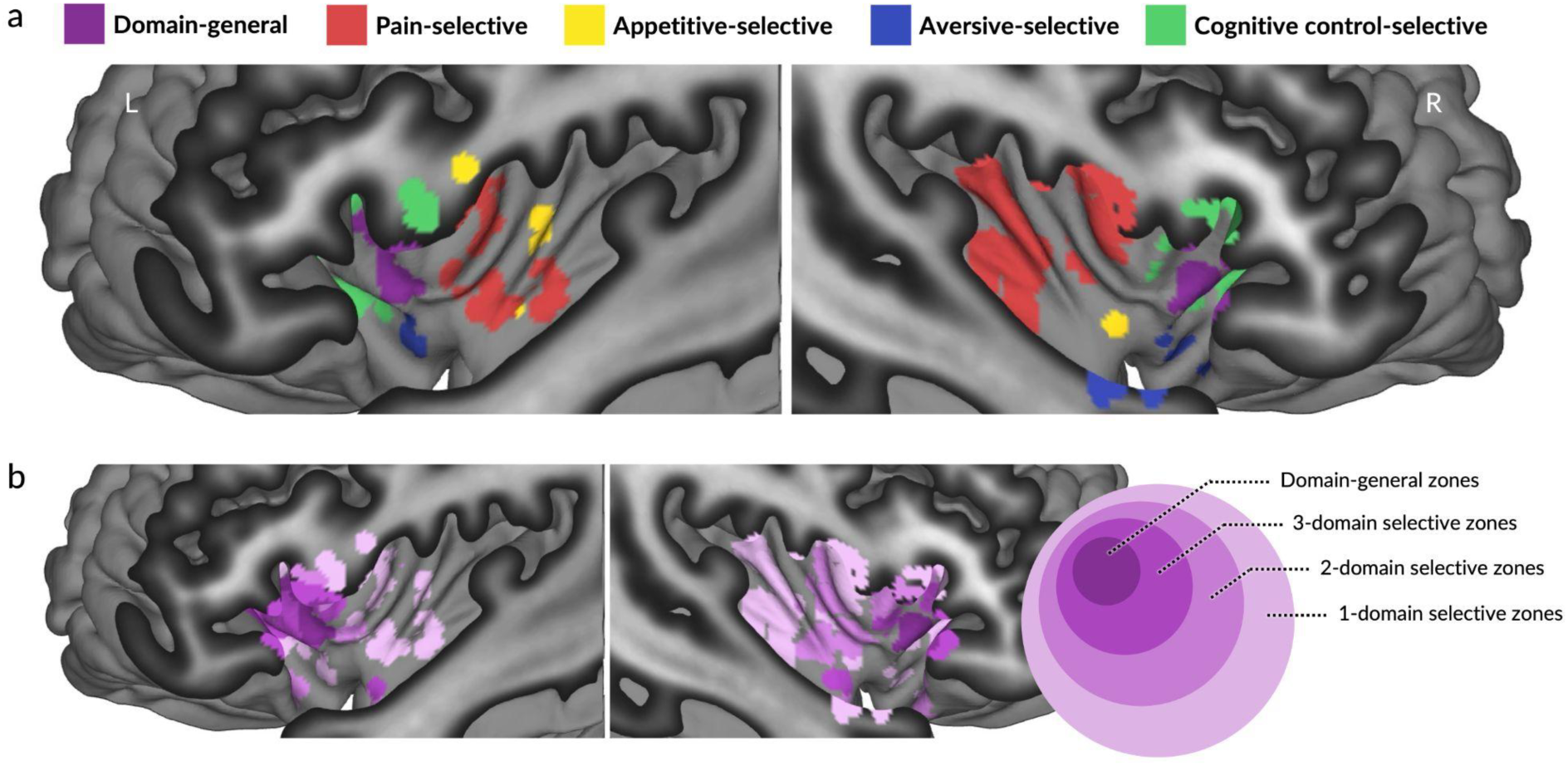
Domain-general and domain-selective zones in the insula. **a,** Domain-general and domain-selective voxels clustered predominantly in specific insular subregions: domain-general voxels (purple) in dorsal anterior insula, pain-selective voxels (red) in mid-posterior insula, appetitive-selective voxels (green) in mid-insula, aversive-selective voxels (blue) in ventral anterior insula, cognitive-selective voxels (yellow) also in dorsal anterior insula, anterior to domain-general voxels. **b,** Spatial gradient of functional convergence in the insula. Color gradient represents the degree of functional convergence, with darker purple indicating higher levels of convergence (i.e., activation across more domains). The map shows a progression from domain-selective zones (lightest purple) in peripheral areas, through zones responsive to two or three domains (intermediate shades of purple), to fully domain-general zones (darkest purple) located centrally in dorsal anterior insula.

We also identified single-domain-selective zones for each domain. Pain-selective zones (320 voxels, red in Fig. 2a) included voxels clustered in mid-posterior insula bilaterally, spanning from middle short to posterior long gyrus. Appetitive-selective zones (75 voxels, yellow) were proximal to pain-selective zones in left dorsal and bilateral ventral mid-insula, spanning anterior and posterior long gyri. Aversive-selective zones (142 voxels, blue) were primarily located in the ventral-most portion of bilateral agranular insula, on anterior inferior gyrus. Cognitive-selective zones (142 voxels, green) were predominantly located in the anterior-most portion of dAIns and most dorsal frontal opercular border bilaterally and spanned anterior and middle short gyri.

These domain-selective zones showed preferential activations for each given domain (Supplementary Fig. 4; see Supplementary Table 1 for further information on each insular zone). These activation patterns were consistent across subdomains and studies with few exceptions.

We further identified zones activated across two or three domains, surrounding domain-general zones activated across all domains (Fig. 2b). Coupled with the findings above, the topography of these zones demonstrated a spatial gradient of convergence, progressing from domain-selective zones (lightest purple) in posterior and ventral insular areas, through zones responsive to two or three domains (intermediate shades of purple), to full multi-domain convergence (darkest purple) in specific dAIns areas. Overall, we identified both domain-general in dAIns and domain-selective zones in surrounding regions, with bilateral symmetry across all domains, revealing a hierarchical progression from specialized processing to multi-domain convergence.

### Functional decoding using Neurosynth topic and term maps

To map the identified insular zones onto psychological topics, we employed meta-analytic functional decoding using Neurosynth^42^. We examined correlations (measured in z-scored point- biserial correlation) with 525 terms and 50 topics derived from 11,406 studies in the Neurosynth database, reporting topics showing higher than standardized correlation 1 and the top 10 most correlated terms per zone (Fig. 3a-b; for a full list of topics, see Supplementary Table 2). These associations provide a data-driven approach to understanding the psychological functions associated with each zone, while acknowledging that topic- and term-based inference is inherently exploratory.

**Fig 3.**
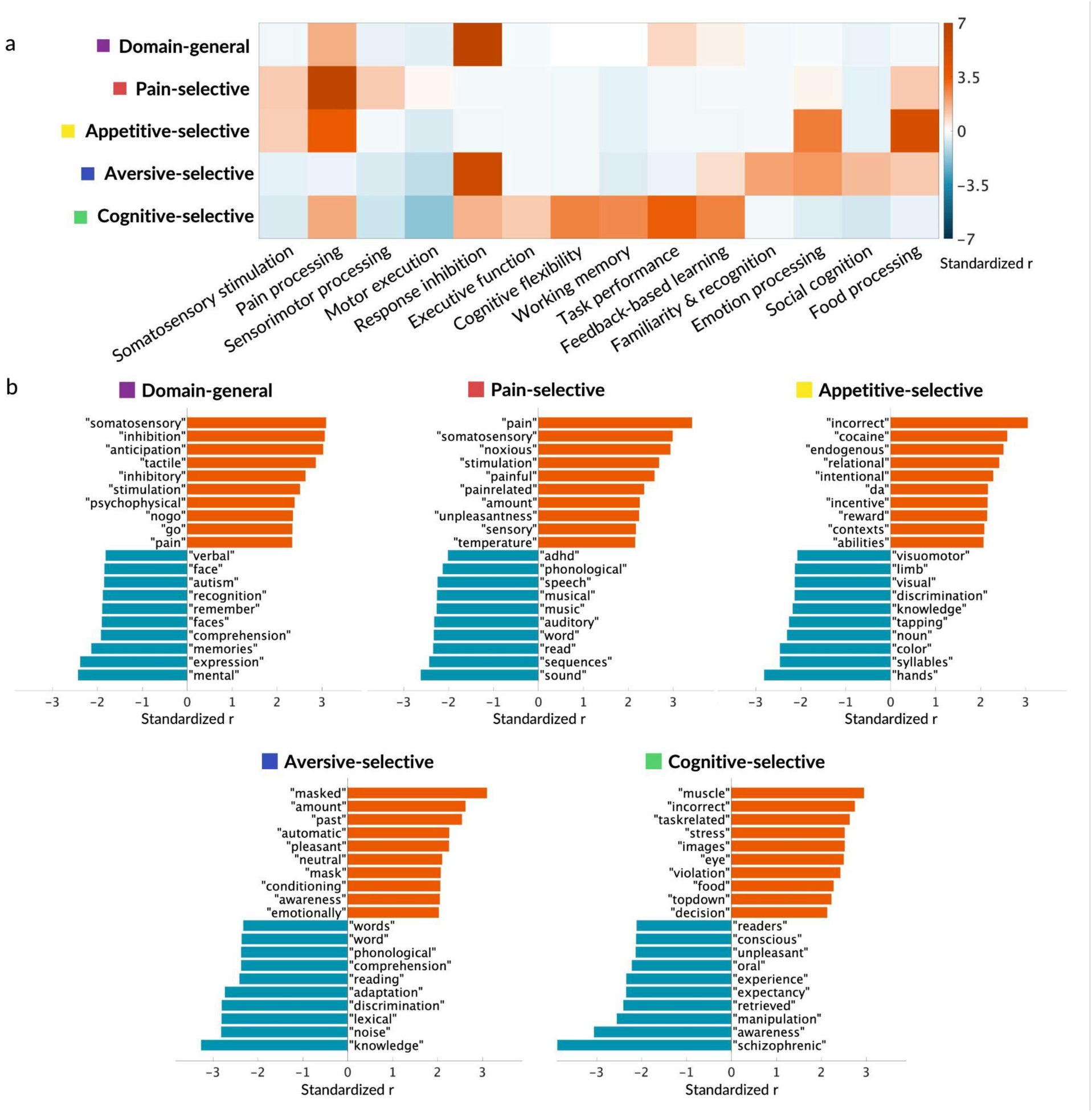
Meta-analytic functional decoding of insular zones using Neurosynth. **a,** Topic- level decoding. Heatmap shows standardized point-biserial correlations between domain- general and domain-selective zones and psychological topics registered in Neurosynth. Topics (x-axis) were selected based on standardized correlation>1 from 50 psychological topics (see Supplementary Fig. 5 for complete topic heatmap). **b,** Term-level decoding. Each subplot shows the top 10 highest and lowest correlating terms for each insular zone, selected from 525 psychological terms. Domain-general zones are highly correlated with a heterogeneous set of topics and terms across all domains, while domain-selective voxels exhibit high correlations with topics and terms related to their designated domains. da: Dopamine

Domain-general zones in dAIns demonstrated high correlations with diverse topics and terms, reflecting their hypothesized functional diversity. The strongest topic associations were with “response inhibition”, “pain processing”, and “task performance”. Supporting this functional heterogeneity, these zones were contained in one of the regions ranked in the top 5% of brain voxels in the analysis of functional diversity using Neurosynth topic maps (uniformity test maps of the same set of 50 topics), showing activation across the most heterogeneous set of topics.

In contrast, domain-selective zones showed function-specific associations matching their designated domains. Pain-selective zones exhibited the strongest topic associations with “pain processing”, “somatosensory stimulation,” and “sensorimotor processing” (but also “food processing”). Among individual terms, the strongest were “pain”, “somatosensory”, “noxious”, and “stimulation”, indicating consistency with pain-related findings in previous literature.

Appetitive-selective zones showed strong associations with topics “food processing”, “pain processing”, and “emotion processing”, and with more appetitive processing-specific terms such as “cocaine”, “incentive”, and “reward”. Aversive-selective zones exhibited the strongest associations with topics “response inhibition”, “emotion processing”, “social cognition”, and terms “masked” (indicating priming), “automatic”, “pleasant”, and “conditioning”, suggesting a close relationship with both preconscious and conscious emotion processing. Cognitive- selective zones showed the strongest associations with topics “task performance”, “cognitive flexibility”, “feedback-based learning”, and “working memory”, and terms “muscle”, “task-related”, “top-down”, and “decision”, indicating a strong link with top-down control of motion and behaviors.

### Coactivation between insular zones and other brain systems

To better understand how these insular zones participate in brain-wide systems, we investigated their coactivation patterns with other regions across 27,072 studies in the Neurosynth database. This approach provides a meta-analytic proxy of functional connectivity^19,66,67^. We also examined the relationship between each extra-insular coactivated system and seven canonical resting-state networks in cortical, subcortical, and cerebellar regions^73–75^.

Domain-general zones showed high coactivation with other putative convergence zones across the brain, including bilateral rostrolateral (rlPFC), right ventrolateral (vlPFC), and right posterior dorsomedial prefrontal cortices (dmPFC), bilateral posterior temporal parietal junction (TPJp), and bilateral anterior midcingulate cortex (aMCC) in the cortex (Fig. 4a). No coactivated voxels were found in subcortical regions or thalamus. These regions largely co-localized with those at the transmodal end of the principal cortical gradient^72^. A majority of voxels coactivating with domain-general zones were located in frontoparietal (32.8%) and default (27.1%) networks. These networks are also among the most transmodal networks across the brain.

**Fig 4.**
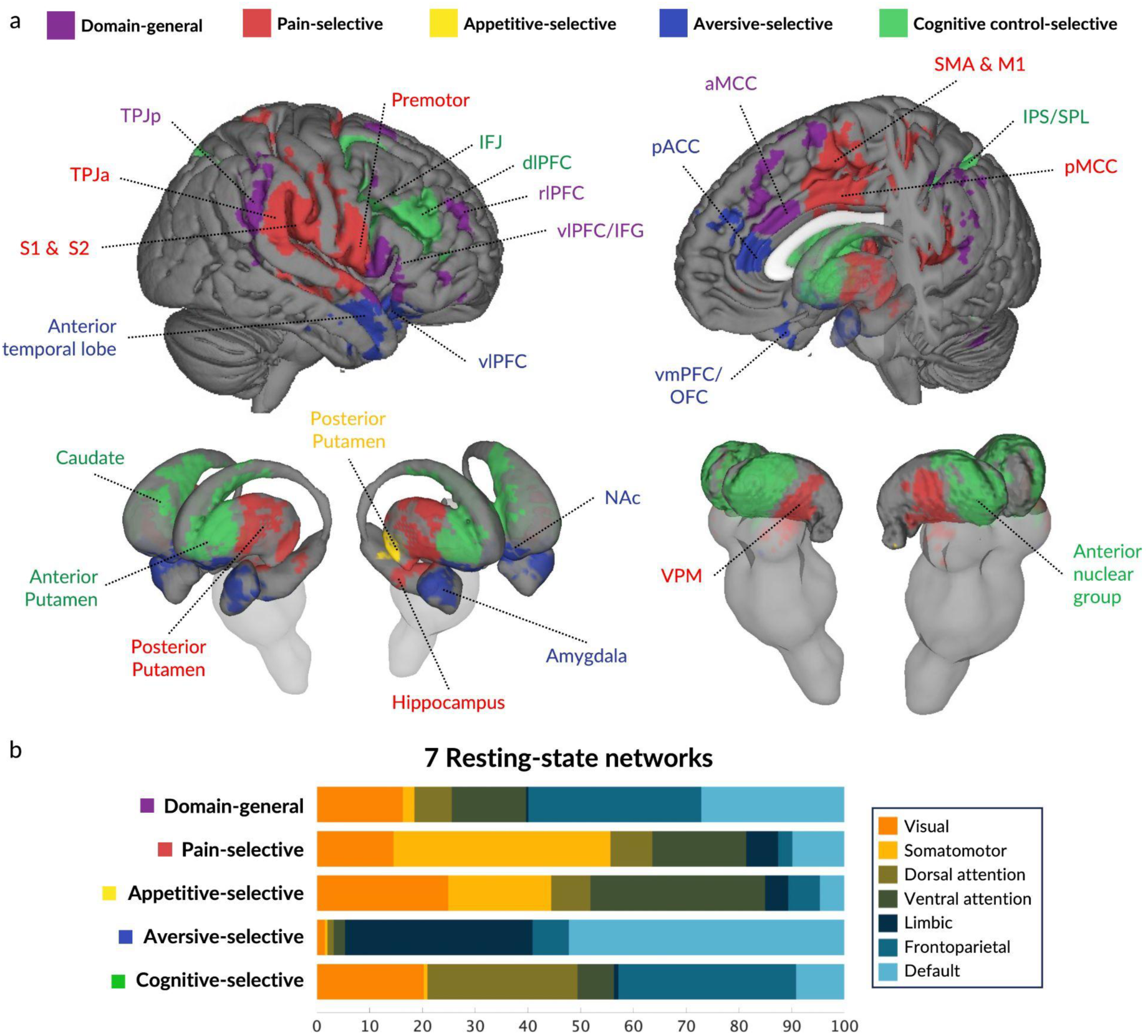
Brain-wide coactivation patterns and resting-state network affiliations of insular zones. **a,** Meta-analytic coactivation patterns between insular zones and extra-insular regions. Coactivated voxels outside of the insula are assigned to the insular zone with which it showed maximum correlation, requiring that maximum correlation to be at least 10% higher than the next highest correlation. aMCC: anterior midcingulate cortex; dlPFC: dorsolateral prefrontal cortex; IFJ: inferior frontal junction; IPS: intraparietal sulcus; SPL: superior parietal lobule; M1: primary motor cortex; NAc: nucleus accumbens; OFC: orbitofrontal cortex; pACC: pregenual anterior cingulate cortex; pMCC: posterior midcingulate cortex; rlPFC: rostrolateral prefrontal cortex; S1: primary somatosensory cortex; S2: secondary somatosensory cortex; SMA: supplementary motor area; TPJa: anterior temporoparietal junction; TPJp: posterior temporoparietal junction; vlPFC: ventrolateral prefrontal cortex; vmPFC: ventromedial prefrontal cortex; VPM: ventral posteromedial nucleus. **b,** Distribution of coactivated regions across seven resting-state networks^73–75^. Bar graph shows the percentage of coactivated voxels overlapping with each network across cortical, subcortical, and brainstem regions.

Pain-selective insular zones showed high coactivation with brain regions involved in somatosensory and somatomotor processing as well as pain processing, including bilateral primary and secondary somatosensory cortices (S1 and S2), right premotor and primary (M1) and supplementary motor area (SMA), bilateral posterior operculum, bilateral anterior TPJ (TPJa), and bilateral posterior medial (pMCC) and dorsal posterior cingulate cortex (dPCC) in the cortex. Subcortical coactivation was observed in ventral posterior thalamus, including ventral posteromedial nucleus (VPM), as well as both posterior and anterior putamen, external and internal globus pallidus, subthalamic nucleus in basal ganglia, and amygdalostriatal transition area and centromedial amygdala. Coactivated voxels were predominantly in somatomotor (41.1%) and ventral attention (17.8%) networks (Fig. 4b). This finding aligns with previous observations in the context of predictive models of pain^76,77^, where a substantial number of pain-predictive voxels were located in somatomotor and ventral attention networks.

Appetitive-selective insular zones showed high coactivation with posterior putamen. These coactivated regions, predominantly located in ventral attention network (33.1%), have been implicated in affective processing across multiple meta-analyses^78^ and individual studies^79,80^.

Aversive-selective insular zones showed the highest coactivation with bilateral orbitofrontal cortex (OFC), anterior medial wall—left ventromedial prefrontal cortex (vmPFC) and right posterior dmPFC, bilateral pregenual anterior cingulate cortices (pACC)—bilateral temporal pole, right superficial amygdala, and right nucleus accumbens (NAc). These regions have been broadly associated with regulation of affective and motivational processes in previous studies. The majority of voxels were found in default mode (52.2%) and limbic (35.5%) networks, with only about 12% in the other networks combined. This provides a striking contrast with somatic pain, which is also aversive but involves very different regions and networks, and helps to elucidate why pain and non-somatic negative affect have been strongly dissociable in prior studies^19,79^.

Cognitive-selective zones showed greater coactivation with bilateral dorsolateral PFC (dlPFC), including inferior frontal junction (IFJ), and bilateral intraparietal sulcus (IPS) and superior parietal lobule (SPL) in the parietal cortex; anterior medial and lateral thalamus across ventral anterior (VA), ventral lateral (VL), lateral dorsal (LD), mediodorsal (MD), ventromedial (VM) nuclei; and bilateral anterior putamen and caudate head. These regions have broadly been associated with cognitive control and goal-based action selection, and a network including dlPFC, IFJ, and anterior insula has been argued to form a neural circuit that selects actions to execute via evidence accumulation^81,82^. In addition, the mid-caudate zone identified here has been selectively associated with executive control in a previous meta-analysis^83^. Coactivating voxels were distributed predominantly among frontoparietal (33.7%) and dorsal attention (28.4%) networks.

### Cytoarchitectonic characterization of domain-general and domain-selective insular zones

To explore the cytoarchitectonic associations of the insular zones, we examined their overlap with the Julich-Brain Cytoarchitectonic Atlas^23,68^, which parcellates the insula into 16 parcels grouped into four clusters (posterior, dorsal anterior, inferior posterior, and ventral anterior) based on cytoarchitectonic characteristics (granular-dysgranular, dysgranular, dysgranular- agranular, and agranular, respectively; Fig. 5). We identified spatial overlap between each insular zone and atlas parcels using Dice Coefficients and selected parcels with coefficients greater than 0.1.

**Fig 5.**
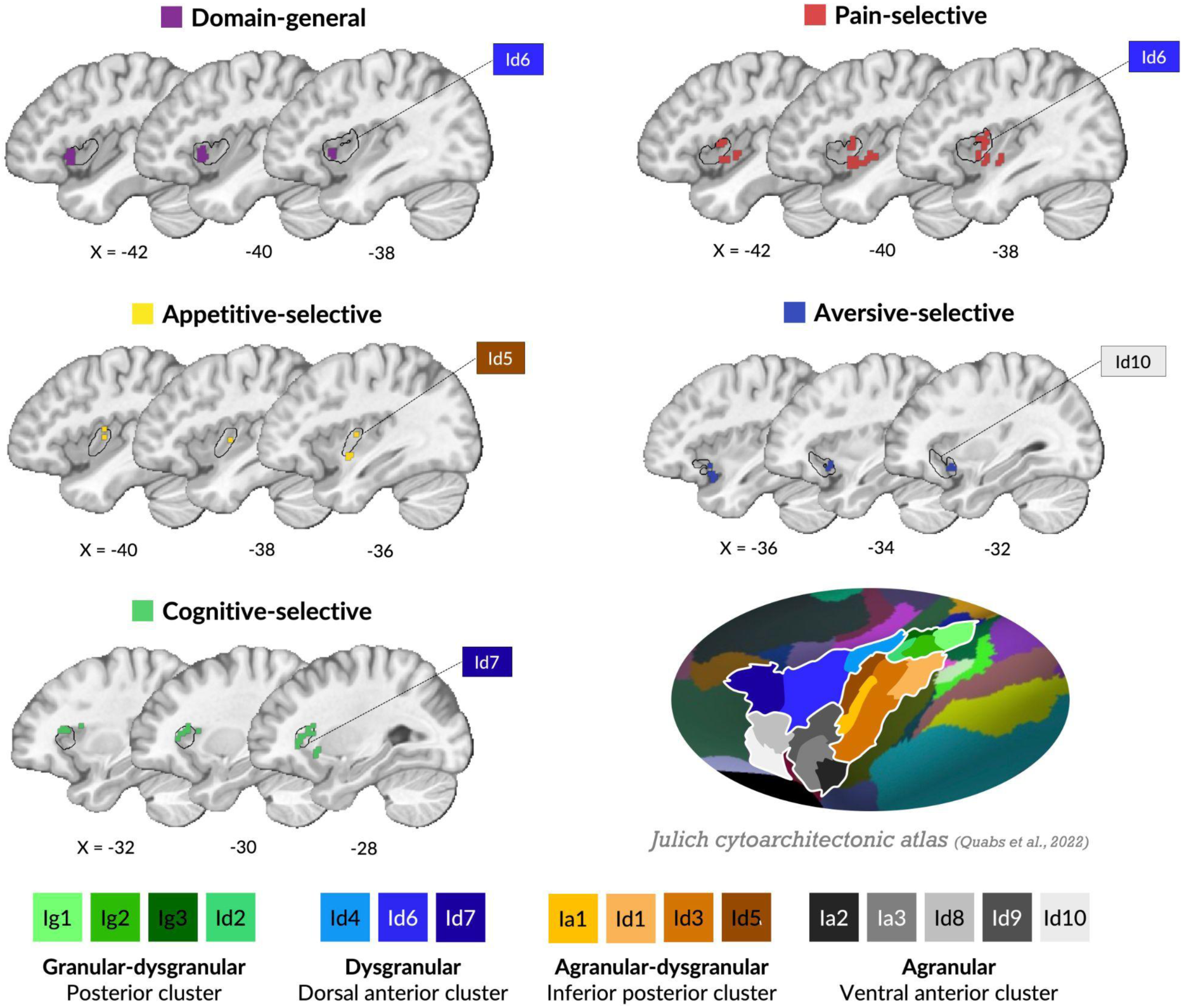
Cytoarchitectonic profiles of insular zones. For each functional zone, only the best matching cytoarchitectonic parcels are shown from the Julich-Brain Cytoarchitectonic Atlas^23,68^ (32 parcels across left and right insula). Inset shows the full cytoarchitectonic atlas of the insula adapted from Quabs et al.^23^ with the authors’ permission.

Identified insular zones mapped onto different cytoarchitectonic regions in the insula, though the overlap was imperfect. Domain-general zones were contained predominantly within left dorsal anterior cluster (dysgranular) and right ventral anterior cluster (agranular), with the strongest overlap with Id6 (see Fig. 5). Cognitive-selective zones also demonstrated high overlap with bilateral dorsal anterior clusters and right ventral anterior cluster, though more dorsal and anterior to domain-general zones, with the strongest overlap in Id7. In contrast, appetitive and aversive-selective zones were primarily in agranular insula, with appetitive-selective zones overlapping most strongly with left inferior posterior cluster (agranular-dysgranular, primarily Id5) and right ventral anterior cluster, while aversive-selective zones overlapped most strongly with bilateral ventral anterior clusters (primarily Id10). Pain-selective zones, anatomically located in mid-posterior insula, showed the strongest overlap with dorsal anterior cluster (primarily Id6).

These zones also extended into inferior posterior (primarily Id5) and posterior (granular-dysgranular; primarily Id3) clusters, spanning across granular, dysgranular, and agranular structures (for a full list of Dice Coefficients, see Supplementary Table 3).

### Neurochemical associations with functional insular zones

To characterize associations with neurotransmitter systems derived from human molecular imaging, we estimated spatial associations (measured in point-biserial correlations) between insular zones and their coactivated brain-wide systems, with PET binding maps of 36 neurotransmitter receptors and transporters across 8 systems from Neuromap^69,70^. We interpreted only associations replicated across at least two independent studies of the same receptor or transporter^84^.

Several neurotransmitter systems showed reliable associations with particular insular zones (Fig. 6a). Mu-opioid (MOR) and cannabinoid (CB1) receptors were strongly associated with domain-general zones and all affective functional domains (appetitive, aversive, and pain-selective zones). Affective domains also had particularly strong associations with serotonin receptor (5-HT1a) and transporter (5HTT) maps, whereas domain-general and cognitive- selective zones showed relatively stronger associations with serotonin 5-HT1b receptor. Both domain-general and pain-selective zones showed strong correlations with glutamate (mGluR5) receptors. Appetitive and cognitive-selective zones shared particularly strong associations with dopamine (D2) receptor distribution. Overall, these findings suggest that different neurotransmitter systems may be differentially involved in specific functional domains, with varying degrees of overlap between the insular zones.

**Fig 6.**
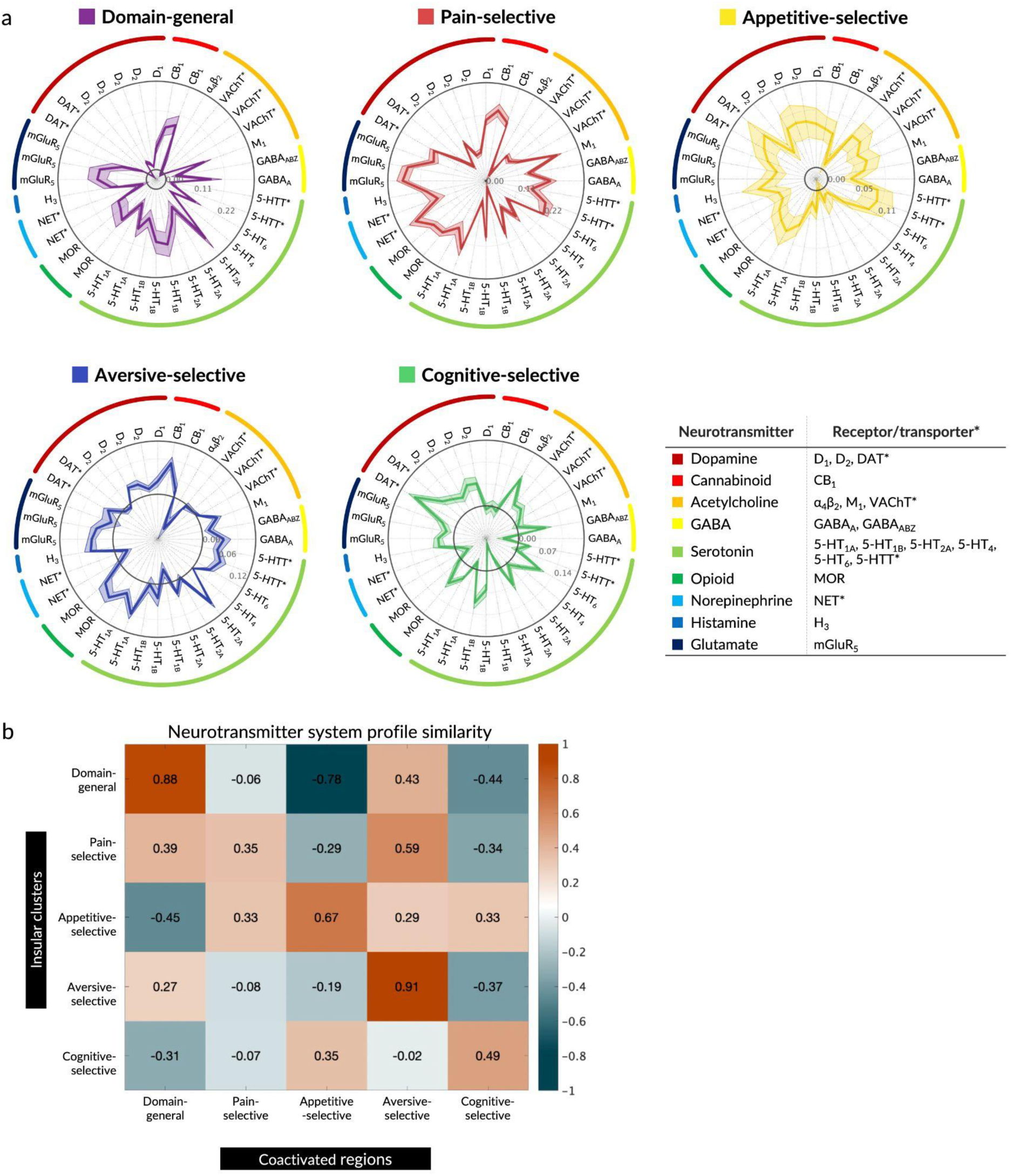
Neurotransmitter system profiles of insular zones. **a,** Spatial similarity between the insular zones and 36 neurotransmitter maps from Neuromap^70,85^ with standard errors estimated through 100 bootstraps. Black rings indicate zero correlation. Inset shows neurotransmitter systems and their corresponding receptors and transporters in the atlas. **b,** Correlation between the insular zones and their extra-insular coactivated systems. Domain-general and selective zones showed highly similar neurotransmitter system profiles to their corresponding extra-insular coactivated regions except pain-selective insular zones and their extra-insular system.

Next, we calculated correlations between the neurotransmitter system profiles of the insular zones and their corresponding coactivated brain-wide networks (Fig. 6b). The neurotransmitter system profiles in domain-general and functionally selective zones in the insula were highly similar to those of their corresponding extra-insular systems (*mean Pearson’s correlation=0.66; range: 0.35–0.91*). For example, the profile of positive spatial associations between domain-general insular zones and CB1, 5-HT1b, mGluR5, and MOR was also present in their coactivated brain-wide systems (e.g., anterior PFC, TPJp, aMCC).

This pattern of neurochemical correspondence was consistent across most domains. Off-target correlations across different functional domains (e.g., between aversive-selective insular zone and appetitive-selective brain-wide systems) were generally weaker than on-target correlations (e.g., between aversive-selective insula and aversive-selective brain-wide systems) and were often negative. This correspondence was strongest for domain-general and aversive domains. Domain-general regions showed on-target similarity *r=0.88* (off-target range: *r=-0.78–0.43)* and aversive processes showed on-target similarity *r=0.91* (off-target range: *r=-0.37–0.27)*. Pain showed the least correspondence, suggesting pain-selective insular zones may have distinct receptor distributions compared to their associated sensorimotor, thalamic, and posterior striatal areas.

## Discussion

The insula contains subregions associated with diverse cognitive, affective, and interoceptive processes, and its hypothesized role in integrating these information streams is thought fundamental for generating unified subjective experiences. However, spatially precise, empirical evidence characterizing functional convergence zones in the insula has been limited. Using large-scale fMRI data of 540 participants systematically sampled across four functional domains and Bayes Factor analyses, we rigorously evaluated both presence and absence of activation across domains. We identified a domain-general convergence zone in bilateral dAIns surrounded by domain-selective zones for pain, appetitive processes, aversive processes, and cognitive control with distinct patterns in their meta-analytic functional profiles, brain-wide coactivation patterns, cytoarchitectonic organization, and neurotransmitter receptor distributions.

Furthermore, inter-domain convergence revealed a structured spatial organization in the insula, progressing from domain-selective zones in peripheral (ventral and posterior) regions to the fully domain-general zones in dAIns, extending beyond previously proposed posterior-to-anterior progression^38,39,86^. These findings support theories that propose hierarchical organization of convergence zones and spatial topography based on functional similarity^36,37^, and align with theories of embodied cognition that emphasize how the progressive integration of bodily signals enables goal-directed behavior and conscious awareness^34,35^.

Our findings identified insular areas of functional convergence and selectivity with increased precision relative to previous work. AIns as heuristically defined by Craig^39^ and dAIns in early meta-analyses^19^ contains both domain-general and functionally selective zones for cognitive control, somatic pain, and/or non-somatic aversive processes in our Bayes Factor analysis. While overlapping BOLD signals alone cannot establish functional convergence, multiple levels of evidence support this result: (a) hierarchical progression from domain-selective to domain-general zones, (b) functional associations based on previous literature, (b) coactivated brain-wide systems, (c) cytoarchitectonic organization, and (d) neurotransmitter system profiles.

Functional associations using Neurosynth meta-analytic topic maps revealed that domain-general zones in dAIns were among the brain regions showing activation across the most heterogeneous set of psychological topics, ranking in the top 5% of all brain voxels for functional diversity, alongside aMCC, IFJ, and medial thalamus. Meta-analytic coactivation analysis further revealed high coactivation with other cortical convergence zones such as rlPFC, TPJp, and aMCC, which are situated at the transmodal end of unimodal-to-transmodal cortical gradients^72^.

Overall coactivation patterns across the identified insular zones showed systematic topographical organization, with organized transitions between functional domains across cortical and subcortical regions. For example, in lateral frontal cortex, we observed an anterior-to-posterior hierarchy from domain-general rlPFC to cognitive control-related dlPFC to pain- related somatomotor regions. This pattern is consistent with theories and findings on prefrontal hierarchy that reflect more abstract, domain-general, and temporally extended processes in anterior regions to more immediate task goals and actions in posterior prefrontal regions^87–90^. In lateral temporal-parietal cortex, domain-general regions in TPJp were situated between cognitive control-related SPL and pain-related TPJa and S2 regions (connected to somatosensory thalamocortical pathways). Similarly, domain-general regions in IFG were positioned between cognitive control-related premotor regions and anterior temporal regions (connected to medial temporal and hippocampal circuits) processing non-somatic aversive responses. In subcortical structures, we found a progression from anterior striatal and thalamic zones specialized for cognitive control to posterior zones specialized for pain, and with appetitive processing potentially localized to the extreme posterior striatum, consistent with areas encoding stable, long-term value in non-human primates^91^.

With respect to canonical resting-state cortical networks, domain-general and domain-selective insular zones and their coactivated systems were unevenly distributed across the networks, but did not align in a one-to-one fashion with them. Instead, the functional systems associated with each insular zone spanned multiple networks. This property is consistent with previous findings that decoding models predicting affective experiences span multiple resting-state networks^92–95^. On one hand, resting-state networks may be related to task-based patterns at a coarse level^96^ and predictive of individual task-evoked topography^97^, but on the other, tasks evoke reorganization of functional connections and networks in ways that cannot be captured by static resting-state patterns^98–100^, including alterations of brain-wide neuronal activity patterns by stimuli and activation of neuromodulatory nuclei^101–105^. This may help explain why the insular zones and coactivated systems we identify here are not reducible to simple combinations of resting-state networks. Instead, activations arise from the bridging of multiple networks.

As with resting-state networks, our functional insular zones mapped onto cytoarchitecture to some degree, but cytoarchitectonic boundaries were not sufficient to describe them completely. Domain-general convergence zones were located largely within cytoarchitectonic region Id6^23^ in dAIns but our results also indicate functional heterogeneity within Id6, including pain and cognitive-selective zones in non-overlapping portions.

The observed cytoarchitectonic mappings were further corroborated by evidence from intracranial studies, which provide more direct neural measurements with higher temporal resolution (electrophysiology) and establish causal links to behavior (direct stimulation).

Domain-general and cognitive-selective zones were predominantly located in dysgranular cortex (Id6 and Id7, respectively), aligning with recent findings suggesting that dysgranular dAIns is a key region for convergence and cognitive control^20,106–112^. Recent intracranial electrophysiology and stimulation studies using the same cytoarchitectonic parcellation further support these associations: while Id7 showed increased activity during task-related salience detection and no responses when stimulated, Id6 showed hypervigilance and anxiety when stimulated—a pattern expected from a multi-modal convergence zone with heightened arousal and vigilance dissociated from specific somatic or affective content^28^. Aversive-selective zones in agranular regions (Id10) align with known limbic system projections^112^ and correspond to regions showing specific responses to negative emotional stimuli in electrophysiological recordings^113^.

Pain-selective zones were a notable exception to this pattern of cytoarchitectonic associations, showing a complex pattern spanning multiple cytoarchitectonic clusters that both aligns with and diverges from previous findings. Beyond Id6’s role in vigilance discussed above, Duong et al.^28^ found that Id6 stimulation evoked visceroception, posterior insular areas Id2 and Ig2 evoked pain, and Id3 evoked somatosensation—findings that align with other stimulation findings on somatic and visceral sensations^6,31,57,114,115^. This broad distribution may reflect both the importance of the insula for interoception and the complex, multidimensional nature of pain, which involves a relatively unique combination of regions across the brain^116,117^. However, our findings diverge from established work showing that posterior granular insula is a key region for processing somatosensory, nociceptive, and pain information^112^. Specifically, we did not observe pain-selective activity in Ig1 (bilateral), Ig2 (left), and Id2 (left) of dorsal posterior insula— regions where many previous studies found pain-selective activity^54,55,118^ and where direct stimulation reliably evokes pain^29,56,119–121^. While right Ig2 and Id2 showed pain-selective activity, they were not the most strongly associated parcels with our pain-selective zones. This apparent discrepancy may reflect the constraints of data harmonization required for analyzing heterogeneous multi-study data. After evaluating several methods, we chose z-scoring for harmonizing data across studies (as detailed in Methods), which affects the detection of pain-selective responses in posterior insula due to pain’s insula-wide activation.

Neurotransmitter receptor and transporter binding patterns from molecular imaging studies in Neuromaps^69,70^ revealed distinct neurochemical patterns across the insular zones. Similar receptor distribution patterns were also observed in brain-wide systems coactivated with these insular zones, suggesting these functionally connected regions share similar capacities for neurochemical modulation. This pattern supports the hypothesis that functional organization is guided by the co-expression of patterns across neurotransmitter systems in functionally related areas, even in the absence of direct structural connections^69,122–125^. This principle is illustrated by recent findings where language-related areas showed similar receptor distribution patterns despite being anatomically distributed^126^, suggesting a common molecular basis for functionally related regions. Furthermore, optogenetic and chemogenetic fMRI studies^127–133^ provide corroborating evidence by demonstrating that manipulating neurotransmitter release from subcortical centers can modulate functional connectivity among regions with shared receptor profiles.

One exception to this pattern of neurochemical correspondence was that the neurotransmitter profile of pain-selective insular zones showed higher similarity to the extra-insular non-somatic negative affect system than to their own extra-insular pain system. This finding implies either pain-related neuromodulatory systems could be more region-specific within the insula, or pain processing in the insula may not be exclusively nociceptive, but may also involve pain-specific affective components, i.e., “pain” representations across nociceptive and vicarious pain, as was recently observed in mid-insula^95^.

These findings highlight the value of our systematic multi-domain sampling approach in revealing both convergent and divergent patterns of functional organization in the insula and present an important direction for future research. This approach recognizes that activity patterns related to broad constructs like “pain”, “cognitive control”, and so forth can only be effectively identified by testing across distinct subdomains with varied stimuli and task designs^134–136^. While large-sample studies^137–141^ enable investigation of how multiple psychological constructs are represented in the brain across large populations, the use of single tasks to represent broad psychological constructs (e.g., an emotional face-matching task to study emotion or an N-back task to study cognitive control and working memory) limits their ability to identify generalizable representations. Characterizing generalizable neural representations requires systematic sampling across multiple implementations of each construct, as demonstrated by our approach through the ANiC and previous work^142^.

Finally, the present study has limitations that should be addressed in future research. First, the impact of specific methodological variables on activation patterns requires further investigation using a larger set of studies with systematically coded methodological variations. Second, while our study systematically sampled four functional domains, several domains important to insular function remain to be examined—e.g., chemosensation such as smell and taste^58,59^, heartbeat perception^51–53,143^, non-pain somatic aversive experiences like breathlessness^144^ and itch^145^, autonomic arousal^143^, inflammation and sickness^25,48–50^, nausea^146^, tussis^147^, and others.

Addressing this limitation will require collaboration and sharing of multiple studies of sufficient size (n ≧ 15). Third, our psychological domain structure represents just one possible ontology for classifying studies^148–151^. The development and validation of ontologies is a complex and important research topic, and there is no single “correct” solution. Future research with extended samples could empirically compare different candidate ontologies, working toward developing psychological categories that are better aligned with the functional architecture of the brain^150,152^. We hope our study inspires future work to expand this approach and address these outstanding long-term goals.

Lastly, our analysis focuses primarily on activation patterns and does not extensively consider information carried by deactivation. While aggregating individual-level contrast images allows for observation of deactivation patterns that can provide valuable insights, interpreting these patterns presents significant challenges, as their underlying mechanisms are less well understood than those of activations. The relationship between BOLD signal decreases and neural activity may vary across brain regions, involving interactions between neural activity, neurovascular coupling, and hemodynamics^153–155^. Beyond this regional variability, BOLD decreases can reflect various neural processes, from inhibition of task-irrelevant activity to resource reallocation or shifts between functional modes^156,157^. Additionally, the choice of baseline condition significantly affects observed deactivation patterns, complicating cross-study comparisons. Given these complexities and the activation-based nature of our subsequent multi-level characterization analyses, we focused our primary Bayes Factor analysis on activation patterns. However, recognizing the potential importance of deactivation, we have included deactivation-based results in Supplementary Fig. 6 for reference. We encourage future studies to further investigate deactivation patterns for various brain functions in and beyond the insula.

Overall, this study provides strong empirical evidence for a fundamental organizing principle in the insula: the coexistence of functionally convergent and selective zones within a brain region long hypothesized to be crucial in integrating diverse information streams. Through systematic sampling across four functional domains and rigorous Bayes Factor analysis of both presence and absence of activation, we reveal a gradient of convergence from domain-selective zones to a multi-domain convergence zone in bilateral dAIns, with an unprecedented combination of spatial coverage, precision, and functional diversity. Our multi-level characterization demonstrates that these functional zones are distinguished by specific patterns of cytoarchitecture, coactivation with other brain regions, and neurotransmitter distribution— providing evidence beyond mere overlap of BOLD signals that these represent distinct functional units. These results reconcile previous work by demonstrating how functional specialization and information convergence coexist within the insula’s hierarchical organization, suggesting a potential role for this architecture in integrating diverse information streams to support unified subjective experiences and the construction of the sense of self.

## Methods

### Study design

This study uses a construct-validation approach, building upon previous research by Kragel et al.^142^ and Van Oudenhove et al.^158^, to explore domain-general and domain-selective representations of four functional domains in the insula: somatic pain, non-somatic appetitive processes, non-somatic aversive processes, and cognitive control. This approach aims to identify brain regions that consistently respond to a particular psychological construct (e.g., pain) across multiple experimental manipulations and studies, ensuring that the voxels identified as domain-general and domain-selective are not driven by specific experimental conditions or study-specific factors, but rather represent the underlying latent construct (for further details on the rationale of this approach, see refs.^142,158^).

Specifically, we systematically sampled participant-level fMRI activation maps from the Affective Neuroimaging Consortium (www.anic.science) database across the four domains above. Each domain includes three subdomains representing different experimental manipulations that engage processes within that domain. Pain domain includes responses to thermal, mechanical, and visceral stimulation; appetitive processes domain includes responses to food, drug, and sexual images; aversive processes domain includes responses to negative images, aversive sounds, and negative social interactions; and cognitive control domain includes responses during working memory, response inhibition, and attention switching tasks. For each subdomain, we included three independent studies using similar experimental manipulations, with 15 randomly sampled participants per study (total *k=36* studies, *n=540*). This hierarchical and balanced design allows us to dissociate insula sub-areas that are selective for a specific domain and those that generalize across all four domains. See Supplementary Table 4 for a full list of studies included in the current dataset.

The current study was a mega-analysis of multiple independent studies. Participants were recruited independently for each study and informed consent was provided by all subjects in accordance with local ethics and institutional review boards. Descriptions of ethics approvals, image acquisition, and demographics are described briefly for all 36 studies in Supplementary Table 4 and in full detail in the corresponding references (see also the Life Sciences Reporting Summary).

### Data harmonization

To ensure compatibility and consistency of the data aggregated from multiple studies, all participant-level activation maps used in the current study were first resampled into a reference space (Study 1^159^) and harmonized using z-scoring normalization across voxels within the insula. By analyzing patterns of values across voxels instead of absolute intensities, this approach helps to account for idiosyncrasies in the scale of activity across studies, which is influenced by various factors such as differences in acquisition, preprocessing, voxel size, contrast weights, and other nuisance factors. This implies that local activity estimates are lower in images with more overall activations across the insula, leading to more conservative results, as domain convergence and selectivity analyses are performed on the relative activation maps. Therefore, we analyzed the standardized means (i.e., mean across insular voxels divided by standard deviation across insular voxels, mean/SD) removed from each activation map during z-scoring normalization at the domain, subdomain, and study levels to assess potential differences in whole-insula activation between domains (see whole-insula activation below). We chose z-scoring after considering several other harmonization methods including ComBat^160^, L2 normalization, and standard deviation normalization. These alternatives were either not applicable due to the nested relationship between scanning parameters and tasks, or did not significantly improve harmonization (see Supplementary Notes for detailed discussion of harmonization approaches and their limitations). Similar normalization procedures are common in multivariate pattern analysis in fMRI and in machine learning and multivariate statistics more broadly (e.g., in profile analysis). Here, this allows us to focus on the relative differences in brain patterns across the four mental constructs while minimizing the impact of study-specific factors.

### Whole-insula activation

To better understand the information removed during z-scoring normalization, we analyzed the standardized means (i.e., mean/SD values removed in normalization) of contrast coefficients across functional domains to check for inter-domain differences in whole-insula activation (see Supplementary Fig. 7). The standardized means of all domains were significantly larger than 0 after FDR correction at *p<0.05*, indicating that all domains activated the insula overall: pain (*mean=0.68,* two-tailed *t(134)=12.74, 95% CI [0.57, 0.78], Cohen’s d = 1.10);* appetitive processes (*mean=0.40, t(134)=5.69, p<0.01, 95% CI [0.26, 0.53], Cohen’s d = 0.49);* aversive processes (*mean=0.13, t(134)=2.56, 95% CI [0.03, 0.24], Cohen’s d = 0.22);* and cognitive control (*mean=0.15, t(134)=2.52, 95% CI [0.03, 0.26], Cohen’s d = 0.22).* Pain showed significantly higher standardized means compared to other domains: versus appetitive processes (*two-tailed t(268)=3.18, 95% CI [0.11, 0.45], Cohen’s d = 0.39),* aversive processes (*t(268)=7.24, 95% CI [0.39, 0.69], Cohen’s d = 0.88),* cognitive control *(t(268)=6.77, 95% CI [0.38, 0.68], Cohen’s d = 0.83, all FDR-corrected p<0.01)*. These results indicate that pain elicits more pronounced and widespread activations across the insula, possibly due to its engagement of a diffuse modulatory system. One consequence is that local posterior insular activity for pain, which was apparent in the pre-normalized maps, was not significant after normalization. This highlights that the main results should be interpreted as relative patterns of activation, with pain showing the strongest insula-wide response.

### Multiclass support vector machine classifier

We trained multi-class linear Support Vector Machine (SVM) classifiers with 5-fold cross-validation for hyperparameter optimization (box constraint). The classifiers were trained to discriminate each domain from the others (one vs. all scheme) using two studies per subdomain for training and a third study as an independent test set for domain and subdomain-level evaluations (e.g., pain from all other domains, thermal pain from all other subdomains). To estimate out-of-study generalization error, we systematically sampled different studies for training and test sets by leaving a different study out in each subdomain for each model, training 100 models in total, and reported the average performance on the test sets (see Supplementary Fig. 8 for a schematic description of study selection).

### Bayes Factors

We calculated Bayes Factors (BFs) at the voxel level using Bayes Factor one-sample t-test with the Jeffrey-Zellner-Siow prior (JZS, Cauchy distribution on effect size; ref.^65^) to identify voxels in the insula exhibiting domain-general or domain-selective activation patterns. BF directly compares the probability of the data under two competing hypotheses: the alternative hypothesis (an effect exists) versus the null hypothesis (no effect).

First, the Bayes Factor for an one-sample t-test is calculated as follows:

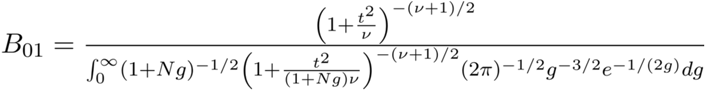

where *t* is the t-statistic, *N* is the sample size, *v* is the degrees of freedom (N - 1), and *r* is the scale factor (set to 0.707, indicating a moderate expected effect size; refs.^65,84^).

We set thresholds at *BF>4.32* for an effect and *BF<0.23* (an inverse of 4.32) for no effect, corresponding to 4.32:1 odds. This decision was supported by previous literature suggesting that Bayes Factors with JZS between 3 and 10 are considered moderate-level evidence^161–163^. These thresholds were chosen to ensure the identification of effects and null effects, minimizing the potential for false positives while maintaining sensitivity to true effects.

Voxels were considered activated only when showing both *BF>4.32* and positive t-statistics. Evidence for no activation included cases of either evidence for no effect (*BF<4.32* with either positive or negative t-statistics) or evidence for deactivation effect (*BF>4.32* with negative t-statistics). By conjunction of BFs across all domains, domain-general voxels required activation across all domains, while domain-selective voxels required activation in their designated domain and evidence for no activation in other domains. Specifically, domain-general voxels were defined as those with *BF>4.32* and *t>0* across all domains. Domain-selective voxels were defined as those with 1) *BF>4.32* and *t>0* for the designated domain, and 2) *BF<0.23* or *BF>4.32* and *t<0* for the other domains.

### Neurosynth functional decoding

We examined spatial correlations between the identified insular zones and two types of meta-analytic maps: 525 term and 50 topic association test maps that show voxels preferentially associated with studies related to particular psychological terms or topics. Topic maps were streamlined from a set of 100 topics (“vs-topics-100”) that were extracted using Latent Dirichlet Allocation (LDA) from abstracts of all articles in the Neurosynth database as of July, 2015 (11,406 articles), after excluding non-psychologically relevant topics. We used standardized point-biserial correlations to calculate spatial correlations between our identified insular zones and these two types of meta-analytic maps for topics and terms. Correlation coefficients were z-scored within each zone to examine the relative strength of correlations across topics for each zone. We report topics showing standardized correlation>1 and the top 10 highest and lowest correlating terms for each insular zone. See Supplementary Table 2 for a full list of Neurosynth topic maps used in the current analysis.

### Neurosynth functional diversity analysis

To assess functional diversity across brain regions, we used the uniformity test maps from the same set of 50 topics, which show voxels consistently activated across studies associated with each topic. For each brain voxel, we calculated the proportion of topics showing significant activation. Voxels were then ranked brain-wide based on this proportion to quantify their functional diversity across psychological topics.

### Coactivation between the insula and other brain areas

We performed a meta-analytic coactivation analysis using activation coordinates from 27,072 studies registered in the Neurosynth repository as of April, 2022. We created meta-analytic binary activation maps using Multi-level kernel density Analysis (MKDA; ref.^164^) with a 4-mm smoothing kernel. In these maps, a voxel was coded as active for a study if the reported peak coordinate was within 4-mm of the given voxel. Using these meta-analytic activation maps, we calculated point-biserial correlations between each domain-general and domain-selective insular zone and each voxel in the cortical and subcortical regions of the whole brain, excluding the insula. The resulting correlations were thresholded at *FDR-corrected q<0.01* and cluster size>100 voxels. Voxels above the threshold were assigned to one of the insular zones based on which zone showed the maximum correlation, provided that the maximum correlation with one of the insular zones was at least 10% higher than the next highest correlation.

To examine the relationship between the coactivated regions and seven resting-state networks (combined from refs.^73–75^), we calculated the percentage of voxels in the coactivated regions that overlapped with each of the seven resting-state networks in the cortical, subcortical, and cerebellar regions.

### Cytoarchitectonic profiling

To determine the best matching cytoarchitectonic parcel for each insular zone, we calculated spatial overlap between each insular zone and each atlas parcel from the Julich-Brain Cytoarchitectonic Atlas^23,68^ (see Fig. 5) using the Dice Coefficient (DC), a measure ranging from 0 (no intersection) to 1 (complete intersection). This atlas includes 16 distinct parcels in the insula, which are regrouped into four clusters (posterior, dorsal anterior, inferior posterior, and ventral anterior) based on their common cytoarchitectonic characteristics (granular-dysgranular, dysgranular, dysgranular-agranular, and agranular, respectively). We selected parcels with a DC higher than 0.1 to report and note which group each insular zone is predominantly associated with.

### Neurotransmitter system profiling

We used a comprehensive receptor and transporter binding map atlas (Neuromap^69,70^) that contains 40 PET-derived binding maps covering 19 neurotransmitter receptors and transports and 9 systems in over 1,200 healthy individuals. The atlas includes maps for the following receptors and transporters (transporters indicated with *): α4β2, M1, VAChT* (acetylcholine); CB1 (cannabinoid); D1, D2, D3, DAT* (dopamine); GABAA/BZ (GABA); mGluR5, NMDA (glutamate); H3 (histamine); NET (norepinephrine); MOR (opioid); 5-HT1a, 5-HT1b, 5-HT2a, 5- HT4, 5-HT6, 5-HTT* (serotonin). Here we used 36 images after removing 4 images due to issues found during implementation.

We calculated point-biserial correlations between each insular zone and each receptor and transporter map in the atlas. Higher correlations indicate the relatively greater density of a given neurotransmitter receptor/transporter. To evaluate the significance and error variability of associations, we bootstrapped each domain-general and selective insular map for 100 iterations. For each iteration, we resampled individual contrast maps and regenerated four insular maps for each domain based on these bootstrap samples. Spatial correlations were then calculated between each bootstrapped insular map and each PET binding map, reporting the mean and standard deviation of these spatial correlations across the 100 bootstrap samples as an estimate of the standard error. For the representativeness of the results, we only interpret the results where 1) there is more than one study on the same neurotransmitter receptor or transporter, and 2) the results from all studies on the same neurotransmitter receptor or transmitter are in agreement. We made exceptions for cases showing meaningful associations based on previous literature. We applied the same approach to the coactivated extra-insular regions of each insular zone. For each insular zone, we computed Pearson’s correlations between its neurotransmitter receptor profile (pattern of receptor densities across different systems) and the receptor profile of its corresponding coactivated regions to test for shared patterns of receptor/transporter distributions.

## References

1. Mesulam MM, Mufson EJ. Insula of the old world monkey. III: Efferent cortical output and comments on function. J Comp Neurol. 1982;212(1):38–52. doi:10.1002/cne.902120104

2. Mufson EJ, Mesulam MM. Insula of the old world monkey. II: Afferent cortical input and comments on the claustrum. J Comp Neurol. 1982;212(1):23–37. doi:10.1002/cne.902120103

3. Augustine JR. Circuitry and functional aspects of the insular lobe in primates including humans. Brain Res Rev. 1996;22(3):229–244. doi:10.1016/S0165-0173(96)00011-2

4. Kurth F, Zilles K, Fox PT, Laird AR, Eickhoff SB. A link between the systems: functional differentiation and integration within the human insula revealed by meta-analysis. Brain Struct Funct. 2010;214(5):519–534. doi:10.1007/s00429-010-0255-z

5. Nieuwenhuys R. Chapter 7 - The insular cortex: A review. In: Hofman MA, Falk D, eds. Progress in Brain Research. Vol 195. Evolution of the Primate Brain. Elsevier; 2012:123-163. doi:10.1016/B978-0-444-53860-4.00007-6

6. Stephani C, Vaca GFB, MacIunas R, Koubeissi MZ, Luders H. Functional neuroanatomy of the insular lobe. Brain Struct Funct. Published online 2011. doi:10.1007/S00429-010-0296-3

7. Türe U, Yaşargil DC, Al-Mefty O, Yaşargil MG. Topographic anatomy of the insular region. J Neurosurg. 1999;90(4):720–733. doi:10.3171/jns.1999.90.4.0720

8. Centanni SW, Janes AC, Haggerty DL, Atwood B, Hopf FW. Better living through understanding the insula: Why subregions can make all the difference. Neuropharmacology. 2021;198:108765. doi:10.1016/j.neuropharm.2021.108765

9. Cerliani L, Thomas RM, Jbabdi S, et al. Probabilistic tractography recovers a rostrocaudal trajectory of connectivity variability in the human insular cortex. Hum Brain Mapp. 2012;33(9):2005–2034. doi:10.1002/hbm.21338

10. Cloutman LL, Binney RJ, Drakesmith M, Parker GJM, Lambon Ralph MA. The variation of function across the human insula mirrors its patterns of structural connectivity: Evidence from in vivo probabilistic tractography. NeuroImage. 2012;59(4):3514–3521. doi:10.1016/j.neuroimage.2011.11.016

11. Ghaziri J, Tucholka A, Girard G, et al. The Corticocortical Structural Connectivity of the Human Insula. Cereb Cortex. 2017;27(2):1216–1228. doi:10.1093/cercor/bhv308

12. Ghaziri J, Tucholka A, Girard G, et al. Subcortical structural connectivity of insular subregions. Sci Rep. 2018;8(1):8596. doi:10.1038/s41598-018-26995-0

13. Jakab A, Molnár PP, Bogner P, Béres M, Berényi EL. Connectivity-based parcellation reveals interhemispheric differences in the insula. Brain Topogr. 2012;25(3):264–271. doi:10.1007/s10548-011-0205-y

14. Mesulam MM, Mufson EJ. The Insula of Reil in Man and Monkey. In: Peters A, Jones EG, eds. Association and Auditory Cortices. Springer US; 1985:179–226. doi:10.1007/978-1-4757-9619-3_5

15. Nomi JS, Schettini E, Broce I, Dick AS, Uddin LQ. Structural Connections of Functionally Defined Human Insular Subdivisions. Cereb Cortex. 2018;28(10):3445–3456. doi:10.1093/cercor/bhx211

16. Deen B, Pitskel NB, Pelphrey KA. Three Systems of Insular Functional Connectivity Identified with Cluster Analysis. Cereb Cortex. 2011;21(7):1498–1506. doi:10.1093/cercor/bhq186

17. Cauda F, Costa T, Torta D, et al. Meta-analytic clustering of the insular cortex. Characterizing the meta-analytic connectivity of the insula when involved in active tasks. NeuroImage. Published online 2012. doi:10.1016/J.NEUROIMAGE.2012.04.012

18. Kelly C, Toro R, Di Martino A, et al. A convergent functional architecture of the insula emerges across imaging modalities. NeuroImage. 2012;61(4):1129–1142. doi:10.1016/j.neuroimage.2012.03.021

19. Chang LJ, Yarkoni T, Khaw MW, Sanfey AG. Decoding the Role of the Insula in Human Cognition: Functional Parcellation and Large-Scale Reverse Inference. Cereb Cortex. 2013;23(3):739–749. doi:10.1093/cercor/bhs065

20. Uddin LQ, Kinnison J, Pessoa L, Anderson ML. Beyond the Tripartite Cognition–Emotion– Interoception Model of the Human Insular Cortex. J Cogn Neurosci. 2014;26(1):16–27. doi:10.1162/jocn_a_00462

21. Nomi JS, Farrant K, Damaraju E, Rachakonda S, Calhoun VD, Uddin LQ. Dynamic functional network connectivity reveals unique and overlapping profiles of insula subdivisions. Hum Brain Mapp. 2016;37(5):1770–1787. doi:10.1002/hbm.23135

22. Tian Y, Zalesky A. Characterizing the functional connectivity diversity of the insula cortex: Subregions, diversity curves and behavior. NeuroImage. 2018;183:716–733. doi:10.1016/j.neuroimage.2018.08.055

23. Quabs J, Caspers S, Schöne C, et al. Cytoarchitecture, probability maps and segregation of the human insula. NeuroImage. 2022;260:119453. doi:10.1016/j.neuroimage.2022.119453

24. Dalley JW, Cardinal RN, Robbins TW. Prefrontal executive and cognitive functions in rodents: neural and neurochemical substrates. Neurosci Biobehav Rev. 2004;28(7):771–784. doi:10.1016/j.neubiorev.2004.09.006

25. Koren T, Yifa R, Amer M, et al. Insular cortex neurons encode and retrieve specific immune responses. Cell. 2021;184(24):5902–5915.e17. doi:10.1016/j.cell.2021.10.013

26. Qadir H, Krimmel SR, Mu C, Poulopoulos A, Seminowicz DA, Mathur BN. Structural Connectivity of the Anterior Cingulate Cortex, Claustrum, and the Anterior Insula of the Mouse. Front Neuroanat. 2018;12. doi:10.3389/fnana.2018.00100

27. Tsai PJ, Keeley RJ, Carmack SA, et al. Converging Structural and Functional Evidence for a Rat Salience Network. Biol Psychiatry. 2020;88(11):867–878. doi:10.1016/j.biopsych.2020.06.023

28. Duong A, Quabs J, Kucyi A, et al. Subjective states induced by intracranial electrical stimulation matches the cytoarchitectonic organization of the human insula. Brain Stimulat. 2023;16(6):1653–1665. doi:10.1016/j.brs.2023.11.001

29. Liberati G, Mulders D, Algoet M, et al. Insular responses to transient painful and non-painful thermal and mechanical spinothalamic stimuli recorded using intracerebral EEG. Sci Rep. 2020;10(1):22319. doi:10.1038/s41598-020-79371-2

30. Mazzola L, Mauguière F, Isnard J. Electrical Stimulations of the Human Insula: Their Contribution to the Ictal Semiology of Insular Seizures. J Clin Neurophysiol Off Publ Am Electroencephalogr Soc. 2017;34(4):307–314. doi:10.1097/WNP.0000000000000382

31. Mazzola L, Mauguière F, Isnard J. Functional mapping of the human insula: Data from electrical stimulations. Rev Neurol (Paris). 2019;175(3):150–156. doi:10.1016/j.neurol.2018.12.003

32. Picard F, Bossaerts P, Bartolomei F. Epilepsy and Ecstatic Experiences: The Role of the Insula. Brain Sci. 2021;11(11):1384. doi:10.3390/brainsci11111384

33. Rachidi I, Minotti L, Martin G, et al. The Insula: A Stimulating Island of the Brain. Brain Sci. 2021;11(11):1533. doi:10.3390/brainsci11111533

34. Barrett LF, Simmons WK. Interoceptive predictions in the brain. Nat Rev Neurosci. 2015;16(7):419–429. doi:10.1038/nrn3950

35. Seth AK. Interoceptive inference, emotion, and the embodied self. Trends Cogn Sci. 2013;17(11):565–573. doi:10.1016/j.tics.2013.09.007

36. Damasio AR. The Brain Binds Entities and Events by Multiregional Activation from Convergence Zones. Neural Comput. 1989;1(1):123–132. doi:10.1162/neco.1989.1.1.123

37. Simmons WK, Barsalou LW. The Similarity-in-Topography Principle: Reconciling Theories of Conceptual Deficits. Cogn Neuropsychol. 2003;20(3-6):451. doi:10.1080/02643290342000032

38. Craig AD. How do you feel? Interoception: the sense of the physiological condition of the body. Nat Rev Neurosci. 2002;3(8):655–666. doi:10.1038/nrn894

39. Craig AD. How do you feel — now? The anterior insula and human awareness. Nat Rev Neurosci. 2009;10(1):59–70. doi:10.1038/nrn2555

40. Varela FJ, Thompson E, Rosch E. The Embodied Mind: Cognitive Science and Human Experience. The MIT Press; 1991:xx, 308.

41. Wilson M. Six views of embodied cognition. Psychon Bull Rev. 2002;9(4):625–636. doi:10.3758/BF03196322

42. Yarkoni T, Poldrack RA, Nichols TE, Van Essen DC, Wager TD. Large-scale automated synthesis of human functional neuroimaging data. Nat Methods. 2011;8(8):665–670. doi:10.1038/nmeth.1635

43. Honey CJ, Thesen T, Donner TH, et al. Slow Cortical Dynamics and the Accumulation of Information over Long Timescales. Neuron. 2012;76(2):423–434. doi:10.1016/j.neuron.2012.08.011

44. Jain S, Vo V, Mahto S, LeBel A, Turek JS, Huth A. Interpretable multi-timescale models for predicting fMRI responses to continuous natural speech. In: Advances in Neural Information Processing Systems. Vol 33. Curran Associates, Inc.; 2020:13738-13749. Accessed October 23, 2024. https://proceedings.neurips.cc/paper/2020/hash/9e9a30b74c49d07d8150c8c83b1ccf07-Abstract.html

45. Menon V, Uddin LQ. Saliency, switching, attention and control: a network model of insula function. Brain Struct Funct. 2010;214(5-6):655–667. doi:10.1007/s00429-010-0262-0

46. Mayer EA, Naliboff BD, Craig ADB. Neuroimaging of the Brain-Gut Axis: From Basic Understanding to Treatment of Functional GI Disorders. Gastroenterology. 2006;131(6):1925–1942. doi:10.1053/j.gastro.2006.10.026

47. Van Oudenhove L, Coen SJ, Aziz Q. Functional brain imaging of gastrointestinal sensation in health and disease. World J Gastroenterol WJG. 2007;13(25):3438–3445. doi:10.3748/wjg.v13.i25.3438

48. Eisenberger NI, Moieni M, Inagaki TK, Muscatell KA, Irwin MR. In Sickness and in Health: The Co-Regulation of Inflammation and Social Behavior. Neuropsychopharmacology. 2017;42(1):242–253. doi:10.1038/npp.2016.141

49. Rolls A. Immunoception: the insular cortex perspective. Cell Mol Immunol. 2023;20(11):1270–1276. doi:10.1038/s41423-023-01051-8

50. Tracey KJ. The inflammatory reflex. Nature. 2002;420(6917):853-859. doi:10.1038/nature01321

51. Critchley HD, Wiens S, Rotshtein P, Öhman A, Dolan RJ. Neural systems supporting interoceptive awareness. Nat Neurosci. 2004;7(2):189–195. doi:10.1038/nn1176

52. Garfinkel SN, Seth AK, Barrett AB, Suzuki K, Critchley HD. Knowing your own heart: Distinguishing interoceptive accuracy from interoceptive awareness. Biol Psychol. 2015;104:65–74. doi:10.1016/j.biopsycho.2014.11.004

53. Hassanpour MS, Simmons WK, Feinstein JS, et al. The Insular Cortex Dynamically Maps Changes in Cardiorespiratory Interoception. Neuropsychopharmacology. 2018;43(2):426–434. doi:10.1038/npp.2017.154

54. Brooks JCW, Tracey I. The insula: A multidimensional integration site for pain. Pain. 2007;128(1):1–2. doi:10.1016/j.pain.2006.12.025

55. Labrakakis C. The Role of the Insular Cortex in Pain. Int J Mol Sci. 2023;24(6):5736. doi:10.3390/ijms24065736

56. Mazzola L, Isnard J, Peyron R, Mauguière F. Stimulation of the human cortex and the experience of pain: Wilder Penfield’s observations revisited. Brain. 2012;135(2):631–640. doi:10.1093/brain/awr265

57. Ostrowsky K, Magnin M, Ryvlin P, Isnard J, Guenot M, Mauguière F. Representation of pain and somatic sensation in the human insula: a study of responses to direct electrical cortical stimulation. Cereb Cortex. Published online 2002. doi:10.1093/CERCOR/12.4.376

58. Veldhuizen MG, Nachtigal D, Teulings L, Gitelman DR, Small DM. The Insular Taste Cortex Contributes to Odor Quality Coding. Front Hum Neurosci. 2010;4. doi:10.3389/fnhum.2010.00058

59. Small DM. Taste representation in the human insula. Brain Struct Funct. 2010;214(5):551–561. doi:10.1007/s00429-010-0266-9

60. Allman JM, Tetreault NA, Hakeem AY, et al. The von Economo neurons in frontoinsular and anterior cingulate cortex in great apes and humans. Brain Struct Funct. Published online 2010. doi:10.1007/S00429-010-0254-0

61. Evrard HC, Forro T, Logothetis NK. Von Economo Neurons in the Anterior Insula of the Macaque Monkey. Neuron. Published online 2012. doi:10.1016/J.NEURON.2012.03.003

62. Hong YW, Yoo Y, Han J, Wager TD, Woo CW. False-positive neuroimaging: Undisclosed flexibility in testing spatial hypotheses allows presenting anything as a replicated finding. NeuroImage. 2019;195:384–395. doi:10.1016/j.neuroimage.2019.03.070

63. Salimi-Khorshidi G, Smith SM, Keltner JR, Wager TD, Nichols TE. Meta-analysis of neuroimaging data: A comparison of image-based and coordinate-based pooling of studies. NeuroImage. 2009;45(3):810–823. doi:10.1016/j.neuroimage.2008.12.039

64. Cronbach LJ, Meehl PE. Construct validity in psychological tests. Psychol Bull. 1955;52(4):281–302. doi:10.1037/h0040957

65. Rouder JN, Speckman PL, Sun D, Morey RD, Iverson G. Bayesian t tests for accepting and rejecting the null hypothesis. Psychon Bull Rev. 2009;16(2):225–237. doi:10.3758/PBR.16.2.225

66. Kober H, Barrett LF, Joseph J, Bliss-Moreau E, Lindquist K, Wager TD. Functional grouping and cortical–subcortical interactions in emotion: A meta-analysis of neuroimaging studies. NeuroImage. 2008;42(2):998–1031. doi:10.1016/j.neuroimage.2008.03.059

67. Laird AR, Eickhoff SB, Rottschy C, Bzdok D, Ray KL, Fox PT. Networks of task co-activations. NeuroImage. 2013;80:505–514. doi:10.1016/j.neuroimage.2013.04.073

68. Amunts K, Mohlberg H, Bludau S, Zilles K. Julich-Brain: A 3D probabilistic atlas of the human brain’s cytoarchitecture. Science. 2020;369(6506):988-992. doi:10.1126/science.abb4588

69. Hansen JY, Shafiei G, Markello RD, et al. Mapping neurotransmitter systems to the structural and functional organization of the human neocortex | Nature Neuroscience. Nat Neurosci. Published online October 27, 2022. doi:10.1038/s41593-022-01186-3

70. Markello RD, Hansen JY, Liu ZQ, et al. neuromaps: structural and functional interpretation of brain maps. Nat Methods. Published online October 6, 2022:1–8. doi:10.1038/s41592-022-01625-w

71. Faillenot I, Heckemann RA, Frot M, Hammers A. Macroanatomy and 3D probabilistic atlas of the human insula. NeuroImage. 2017;150:88–98. doi:10.1016/j.neuroimage.2017.01.073

72. Margulies DS, Ghosh SS, Goulas A, et al. Situating the default-mode network along a principal gradient of macroscale cortical organization. Proc Natl Acad Sci. 2016;113(44):12574–12579. doi:10.1073/pnas.1608282113

73. Buckner RL, Krienen FM, Castellanos A, Diaz JC, Yeo BTT. The organization of the human cerebellum estimated by intrinsic functional connectivity. J Neurophysiol. 2011;106(5):2322–2345. doi:10.1152/jn.00339.2011

74. Choi EY, Yeo BTT, Buckner RL. The organization of the human striatum estimated by intrinsic functional connectivity. J Neurophysiol. 2012;108(8):2242–2263. doi:10.1152/jn.00270.2012

75. Yeo BT, Krienen FM, Sepulcre J, et al. The organization of the human cerebral cortex estimated by intrinsic functional connectivity. J Neurophysiol. 2011;106(3):1125–1165. doi:10.1152/jn.00338.2011

76. Kragel P, Wager TD. Reproducible, Generalizable Brain Models of Affective Processes. In: Neta M, Haas IJ, eds. Emotion in the Mind and Body. Nebraska Symposium on Motivation. Springer International Publishing; 2019:221–263. doi:10.1007/978-3-030-27473-3_8

77. Wager TD, Atlas LY, Lindquist MA, Roy M, Woo CW, Kross E. An fMRI-Based Neurologic Signature of Physical Pain. N Engl J Med. 2013;368(15):1388–1397. doi:10.1056/NEJMoa1204471

78. Satpute AB, Kang J, Bickart KC, Yardley H, Wager TD, Barrett LF. Involvement of Sensory Regions in Affective Experience: A Meta-Analysis. Front Psychol. 2015;6. doi:10.3389/fpsyg.2015.01860

79. Čeko M, Kragel PA, Woo CW, López-Solà M, Wager TD. Common and stimulus-type-specific brain representations of negative affect. Nat Neurosci. 2022;25(6):760–770. doi:10.1038/s41593-022-01082-w

80. Kragel PA, Reddan MC, LaBar KS, Wager TD. Emotion schemas are embedded in the human visual system. Sci Adv. Published online 2019:16.

81. Gluth S, Rieskamp J, Büchel C. Deciding when to decide: time-variant sequential sampling models explain the emergence of value-based decisions in the human brain. J Neurosci Off J Soc Neurosci. 2012;32(31):10686–10698. doi:10.1523/JNEUROSCI.0727-12.2012

82. Hutcherson CA, Tusche A. Evidence accumulation, not ‘self-control’, explains dorsolateral prefrontal activation during normative choice. Kable JW, Frank MJ, eds. eLife. 2022;11:e65661. doi:10.7554/eLife.65661

83. Pauli WM, O’Reilly RC, Yarkoni T, Wager TD. Regional specialization within the human striatum for diverse psychological functions. Proc Natl Acad Sci. 2016;113(7):1907–1912. doi:10.1073/pnas.1507610113

84. Bo K, Kraynak TE, Kwon M, Sun M, Gianaros PJ, Wager TD. A systems identification approach using Bayes factors to deconstruct the brain bases of emotion regulation. Nat Neurosci. Published online March 22, 2024:1–13. doi:10.1038/s41593-024-01605-7

85. Hansen JY, Markello RD, Vogel JW, Seidlitz J, Bzdok D, Misic B. Mapping gene transcription and neurocognition across human neocortex. Nat Hum Behav. 2021;5(9):1240–1250. doi:10.1038/s41562-021-01082-z

86. Kuehn E, Mueller K, Lohmann G, Schuetz-Bosbach S. Interoceptive awareness changes the posterior insula functional connectivity profile. Brain Struct Funct. 2016;221(3):1555–1571. doi:10.1007/s00429-015-0989-8

87. Badre D, D’Esposito M. Functional Magnetic Resonance Imaging Evidence for a Hierarchical Organization of the Prefrontal Cortex. J Cogn Neurosci. 2007;19(12):2082–2099. doi:10.1162/jocn.2007.19.12.2082

88. Christoff K, Keramatian K, Gordon AM, Smith R, Mädler B. Prefrontal organization of cognitive control according to levels of abstraction. Brain Res. 2009;1286:94–105. doi:10.1016/j.brainres.2009.05.096

89. Dixon ML, Fox KCR, Christoff K. Evidence for rostro-caudal functional organization in multiple brain areas related to goal-directed behavior. Brain Res. 2014;1572:26–39. doi:10.1016/j.brainres.2014.05.012

90. Koechlin E, Ody C, Kouneiher F. The Architecture of Cognitive Control in the Human Prefrontal Cortex. Science. 2003;302(5648):1181-1185. doi:10.1126/science.1088545

91. Kim HF, Hikosaka O. Distinct Basal Ganglia Circuits Controlling Behaviors Guided by Flexible and Stable Values. Neuron. 2013;79(5):1001–1010. doi:10.1016/j.neuron.2013.06.044

92. Chang LJ, Gianaros PJ, Manuck SB, Krishnan A, Wager TD. A Sensitive and Specific Neural Signature for Picture-Induced Negative Affect. Adolphs R, ed. PLOS Biol. 2015;13(6):e1002180. doi:10.1371/journal.pbio.1002180

93. Kragel PA, Koban L, Barrett LF, Wager TD. Representation, Pattern Information, and Brain Signatures: From Neurons to Neuroimaging. Neuron. 2018;99(2):257–273. doi:10.1016/j.neuron.2018.06.009

94. van’t Hof SR, Van Oudenhove L, Janssen E, et al. The Brain Activation-Based Sexual Image Classifier (BASIC): A Sensitive and Specific fMRI Activity Pattern for Sexual Image Processing. Cereb Cortex N Y N 1991. 2022;32(14):3014-3030. doi:10.1093/cercor/bhab397

95. Zhou F, Zhao W, Qi Z, et al. A distributed fMRI-based signature for the subjective experience of fear. Nat Commun. 2021;12(1):6643. doi:10.1038/s41467-021-26977-3

96. Smith SM, Fox PT, Miller KL, et al. Correspondence of the brain’s functional architecture during activation and rest. Proc Natl Acad Sci. 2009;106(31):13040–13045. doi:10.1073/pnas.0905267106

97. Tavor I, Jones OP, Mars RB, Smith SM, Behrens TE, Jbabdi S. Task-free MRI predicts individual differences in brain activity during task performance. Science. 2016;352(6282):216-220. doi:10.1126/science.aad8127

98. Finn ES. Is it time to put rest to rest? Trends Cogn Sci. 2021;25(12):1021–1032. doi:10.1016/j.tics.2021.09.005

99. Greene AS, Gao S, Noble S, Scheinost D, Constable RT. How Tasks Change Whole-Brain Functional Organization to Reveal Brain-Phenotype Relationships. Cell Rep. 2020;32(8). doi:10.1016/j.celrep.2020.108066

100. Zheng W, Woo CW, Yao Z, et al. Pain-Evoked Reorganization in Functional Brain Networks. Cereb Cortex. 2020;30(5):2804–2822. doi:10.1093/cercor/bhz276

101. Allen WE, Chen MZ, Pichamoorthy N, et al. Thirst regulates motivated behavior through modulation of brainwide neural population dynamics. Science. 2019;364(6437):eaav3932. doi:10.1126/science.aav3932

102. Briggs F, Mangun GR, Usrey WM. Attention enhances synaptic efficacy and the signal-to-noise ratio in neural circuits. Nature. 2013;499(7459):476-480. doi:10.1038/nature12276

103. Jung WB, Jiang H, Lee S, Kim SG. Dissection of brain-wide resting-state and functional somatosensory circuits by fMRI with optogenetic silencing. Proc Natl Acad Sci. 2022;119(4):e2113313119. doi:10.1073/pnas.2113313119

104. Leong ATL, Chan RW, Gao PP, et al. Long-range projections coordinate distributed brain-wide neural activity with a specific spatiotemporal profile. Proc Natl Acad Sci. 2016;113(51):E8306–E8315. doi:10.1073/pnas.1616361113

105. Tu W, Ma Z, Zhang N. Brain network reorganization after targeted attack at a hub region. NeuroImage. 2021;237:118219. doi:10.1016/j.neuroimage.2021.118219

106. Adamic EM, Teed AR, Avery JA, Cruz F de la, Khalsa SS. Hemispheric divergence of interoceptive processing across psychiatric disorders. eLife. 2024;13. doi:10.7554/eLife.92820.2

107. Adolfi F, Couto B, Richter F, et al. Convergence of interoception, emotion, and social cognition: A twofold fMRI meta-analysis and lesion approach. Cortex. 2017;88:124–142. doi:10.1016/j.cortex.2016.12.019

108. Bastuji H, Frot M, Perchet C, Hagiwara K, Garcia-Larrea L. Convergence of sensory and limbic noxious input into the anterior insula and the emergence of pain from nociception. Sci Rep. 2018;8(1):13360. doi:10.1038/s41598-018-31781-z

109. Evrard HC. The Organization of the Primate Insular Cortex. Front Neuroanat. 2019;13. Accessed August 18, 2022. https://www.frontiersin.org/articles/10.3389/fnana.2019.00043

110. Garfinkel SN, Critchley HD. Interoception, emotion and brain: new insights link internal physiology to social behaviour. Commentary on: Soc Cogn Affect Neurosci. 2013;8(3):231–234. doi:10.1093/scan/nss140

111. Krockenberger MS, Saleh-Mattesich TO, Evrard HC. Cytoarchitectonic and connection stripes in the dysgranular insular cortex in the macaque monkey. J Comp Neurol. 2023;531(18):2019–2043. doi:10.1002/cne.25571

112. Uddin LQ, Nomi JS, Hebert-Seropian B, Ghaziri J, Boucher O. Structure and function of the human insula. J Clin Neurophysiol Off Publ Am Electroencephalogr Soc. 2017;34(4):300–306. doi:10.1097/WNP.0000000000000377

113. Yih J, Beam DE, Fox KCR, Parvizi J. Intensity of affective experience is modulated by magnitude of intracranial electrical stimulation in human orbitofrontal, cingulate and insular cortices. Soc Cogn Affect Neurosci. 2019;14(4):339–351. doi:10.1093/scan/nsz015

114. Afif A, Minotti L, Kahane P, Hoffmann D. Anatomofunctional organization of the insular cortex: A study using intracerebral electrical stimulation in epileptic patients. Epilepsia. Published online 2010. doi:10.1111/J.1528-1167.2010.02755.X

115. Soulier H, Mauguière F, Catenoix H, et al. Visceral and emotional responses to direct electrical stimulations of the cortex. Ann Clin Transl Neurol. 2023;10(1):5–17. doi:10.1002/acn3.51694

116. Woo CW, Koban L, Kross E, et al. Separate neural representations for physical pain and social rejection. Nat Commun. 2014;5(1):5380. doi:10.1038/ncomms6380

117. Woo CW, Schmidt L, Krishnan A, et al. Quantifying cerebral contributions to pain beyond nociception. Nat Commun. 2017;8(1):14211. doi:10.1038/ncomms14211

118. Segerdahl AR, Mezue M, Okell TW, Farrar JT, Tracey I. The dorsal posterior insula subserves a fundamental role in human pain. Nat Neurosci. Published online 2015. doi:10.1038/NN.3969

119. Frot M, Magnin M, Mauguière F, Garcia-Larrea L. Human SII and Posterior Insula Differently Encode Thermal Laser Stimuli. Cereb Cortex. 2007;17(3):610–620. doi:10.1093/cercor/bhk007

120. Mazzola L, Isnard J, Mauguière F. Somatosensory and pain responses to stimulation of the second somatosensory area (SII) in humans. A comparison with SI and insular responses. Cereb Cortex. Published online 2006. doi:10.1093/CERCOR/BHJ038

121. Mazzola L, Isnard J, Peyron R, Guénot M, Mauguière F. Somatotopic organization of pain responses to direct electrical stimulation of the human insular cortex. Pain. Published online 2009. doi:10.1016/J.PAIN.2009.07.014

122. Goulas A, Changeux JP, Wagstyl K, Amunts K, Palomero-Gallagher N, Hilgetag CC. The natural axis of transmitter receptor distribution in the human cerebral cortex. Proc Natl Acad Sci. 2021;118(3):e2020574118. doi:10.1073/pnas.2020574118

123. Palomero-Gallagher N, Zilles K. Cortical layers: Cyto-, myelo-, receptor- and synaptic architecture in human cortical areas. NeuroImage. 2019;197:716–741. doi:10.1016/j.neuroimage.2017.08.035

124. Zachlod D, Palomero-Gallagher N, Dickscheid T, Amunts K. Mapping Cytoarchitectonics and Receptor Architectonics to Understand Brain Function and Connectivity. Biol Psychiatry. 2023;93(5):471–479. doi:10.1016/j.biopsych.2022.09.014

125. Suárez LE, Markello RD, Betzel RF, Misic B. Linking Structure and Function in Macroscale Brain Networks. Trends Cogn Sci. 2020;24(4):302–315. doi:10.1016/j.tics.2020.01.008

126. Zilles K, Bacha-Trams M, Palomero-Gallagher N, Amunts K, Friederici AD. Common molecular basis of the sentence comprehension network revealed by neurotransmitter receptor fingerprints. Cortex. 2015;63:79–89. doi:10.1016/j.cortex.2014.07.007

127. Ferenczi EA, Zalocusky KA, Liston C, et al. Prefrontal cortical regulation of brainwide circuit dynamics and reward-related behavior. Science. 2016;351(6268):aac9698. doi:10.1126/science.aac9698

128. Giorgi A, Migliarini S, Galbusera A, et al. Brain-wide Mapping of Endogenous Serotonergic Transmission via Chemogenetic fMRI. Cell Rep. 2017;21(4):910–918. doi:10.1016/j.celrep.2017.09.087

129. Grimm C, Duss SN, Privitera M, et al. Locus Coeruleus firing patterns selectively modulate brain activity and dynamics. Published online August 29, 2022:2022.08.29.505672. doi:10.1101/2022.08.29.505672

130. Hamada HT, Abe Y, Takata N, Taira M, Tanaka KF, Doya K. Optogenetic activation of dorsal raphe serotonin neurons induces brain-wide activation. Nat Commun. 2024;15(1):4152. doi:10.1038/s41467-024-48489-6

131. Ioanas HI, Saab BJ, Rudin M. Whole-brain opto-fMRI map of mouse VTA dopaminergic activation reflects structural projections with small but significant deviations. Transl Psychiatry. 2022;12(1):1–10. doi:10.1038/s41398-022-01812-5

132. Lohani S, Poplawsky AJ, Kim SG, Moghaddam B. Unexpected global impact of VTA dopamine neuron activation as measured by opto-fMRI. Mol Psychiatry. 2017;22(4):585–594. doi:10.1038/mp.2016.102

133. Turchi J, Chang C, Ye FQ, et al. The Basal Forebrain Regulates Global Resting-State fMRI Fluctuations. Neuron. 2018;97(4):940–952.e4. doi:10.1016/j.neuron.2018.01.032

134. Gustavson DE, Stallings MC, Corley RP, Miyake A, Hewitt JK, Friedman NP. Executive functions and substance use: Relations in late adolescence and early adulthood. J Abnorm Psychol. 2017;126(2):257–270. doi:10.1037/abn0000250

135. Miyake A, Friedman NP, Rettinger DA, Shah P, Hegarty M. How are visuospatial working memory, executive functioning, and spatial abilities related? A latent-variable analysis. J Exp Psychol Gen. 2001;130(4):621–640. doi:10.1037//0096-3445.130.4.621

136. Reineberg AE, Banich MT, Wager TD, Friedman NP. Context-specific activations are a hallmark of the neural basis of individual differences in general executive function. NeuroImage. 2022;249:118845. doi:10.1016/j.neuroimage.2021.118845

137. Casey BJ, Cannonier T, Conley MI, et al. The Adolescent Brain Cognitive Development (ABCD) study: Imaging acquisition across 21 sites. Dev Cogn Neurosci. 2018;32:43–54. doi:10.1016/j.dcn.2018.03.001

138. Van Essen DC, Ugurbil K, Auerbach E, et al. The Human Connectome Project: A data acquisition perspective. NeuroImage. 2012;62(4):2222–2231. doi:10.1016/j.neuroimage.2012.02.018

139. Miller KL, Alfaro-Almagro F, Bangerter NK, et al. Multimodal population brain imaging in the UK Biobank prospective epidemiological study. Nat Neurosci. 2016;19(11):1523–1536. doi:10.1038/nn.4393

140. Shafto MA, Tyler LK, Dixon M, et al. The Cambridge Centre for Ageing and Neuroscience (Cam-CAN) study protocol: a cross-sectional, lifespan, multidisciplinary examination of healthy cognitive ageing. BMC Neurol. 2014;14:204. doi:10.1186/s12883-014-0204-1

141. Berardi G, Frey-Law L, Sluka KA, et al. Multi-Site Observational Study to Assess Biomarkers for Susceptibility or Resilience to Chronic Pain: The Acute to Chronic Pain Signatures (A2CPS) Study Protocol. Front Med. 2022;9. doi:10.3389/fmed.2022.849214

142. Kragel PA, Kano M, Van Oudenhove L, et al. Generalizable representations of pain, cognitive control, and negative emotion in medial frontal cortex. Nat Neurosci. 2018;21(2):283–289. doi:10.1038/s41593-017-0051-7

143. Hassanpour MS, Yan L, Wang DJJ, et al. How the heart speaks to the brain: neural activity during cardiorespiratory interoceptive stimulation. Philos Trans R Soc B Biol Sci. 2016;371(1708):20160017. doi:10.1098/rstb.2016.0017

144. Faull OK, Hayen A, Pattinson KTS. Breathlessness and the body: Neuroimaging clues for the inferential leap. Cortex. 2017;95:211–221. doi:10.1016/j.cortex.2017.07.019

145. Ross SE. Pain and itch: insights into the neural circuits of aversive somatosensation in health and disease. Curr Opin Neurobiol. 2011;21(6):880–887. doi:10.1016/j.conb.2011.10.012

146. Napadow V, Sheehan JD, Kim J, et al. The Brain Circuitry Underlying the Temporal Evolution of Nausea in Humans. Cereb Cortex N Y NY. 2013;23(4):806–813. doi:10.1093/cercor/bhs073

147. Farrell MJ, Mazzone SB. Sensations and regional brain responses evoked by tussive stimulation of the airways. Respir Physiol Neurobiol. 2014;204:58–63. doi:10.1016/j.resp.2014.06.009

148. Bolt T, Nomi JS, Arens R, et al. Ontological Dimensions of Cognitive-Neural Mappings. Neuroinformatics. 2020;18(3):451–463. doi:10.1007/s12021-020-09454-y

149. Insel T, Cuthbert B, Garvey M, et al. Research Domain Criteria (RDoC): Toward a New Classification Framework for Research on Mental Disorders. Am J Psychiatry. 2010;167(7):748–751. doi:10.1176/appi.ajp.2010.09091379

150. Poldrack RA, Yarkoni T. From Brain Maps to Cognitive Ontologies: Informatics and the Search for Mental Structure. Annu Rev Psychol. 2016;67(1):587–612. doi:10.1146/annurev-psych-122414-033729

151. Turner JA, Laird AR. The Cognitive Paradigm Ontology: Design and Application. Neuroinformatics. 2012;10(1):57–66. doi:10.1007/s12021-011-9126-x

152. Lindquist KA, Wager TD, Kober H, Bliss-Moreau E, Barrett LF. The brain basis of emotion: A meta-analytic review. Behav Brain Sci. 2012;35(3):121–143. doi:10.1017/S0140525X11000446

153. Hayes DJ, Huxtable AG. Interpreting Deactivations in Neuroimaging. Front Psychol. 2012;3:27. doi:10.3389/fpsyg.2012.00027

154. Frankenstein U, Wennerberg A, Richter W, et al. Activation and deactivation in blood oxygenation level dependent functional magnetic resonance imaging. Concepts Magn Reson Part A. 2003;16A(1):63-70. doi:10.1002/cmr.a.10054

155. Mishra AM, Ellens DJ, Schridde U, et al. Where fMRI and Electrophysiology Agree to Disagree: Corticothalamic and Striatal Activity Patterns in the WAG/Rij Rat. J Neurosci. 2011;31(42):15053–15064. doi:10.1523/JNEUROSCI.0101-11.2011

156. Viviani R, Dommes L, Bosch JE, Labek K. Segregation, connectivity, and gradients of deactivation in neural correlates of evidence in social decision making. NeuroImage. 2020;223:117339. doi:10.1016/j.neuroimage.2020.117339

157. Hasson U, Nusbaum HC, Small SL. Task-dependent organization of brain regions active during rest. Proc Natl Acad Sci. 2009;106(26):10841–10846. doi:10.1073/pnas.0903253106

158. Van Oudenhove L, Kragel PA, Dupont P, et al. Common and distinct neural representations of aversive somatic and visceral stimulation in healthy individuals. Nat Commun. 2020;11(1):5939. doi:10.1038/s41467-020-19688-8

159. Atlas LY, Bolger N, Lindquist MA, Wager TD. Brain Mediators of Predictive Cue Effects on Perceived Pain. J Neurosci. 2010;30(39):12964–12977. doi:10.1523/JNEUROSCI.0057-10.2010

160. Johnson WE, Li C, Rabinovic A. Adjusting batch effects in microarray expression data using empirical Bayes methods. Biostatistics. 2007;8(1):118–127. doi:10.1093/biostatistics/kxj037

161. Jeffreys H. Theory of Probability. Clarendon Press; 1939.

162. van Doorn J, van den Bergh D, Böhm U, et al. The JASP guidelines for conducting and reporting a Bayesian analysis. Psychon Bull Rev. 2021;28(3):813–826. doi:10.3758/s13423-020-01798-5

163. Kass RE, Raftery AE. Bayes Factors. J Am Stat Assoc. 1995;90(430):773-795. doi:10.1080/01621459.1995.10476572

164. Kober H, Wager TD. Meta-analysis of neuroimaging data. WIREs Cogn Sci. 2010;1(2):293–300. doi:10.1002/wcs.41

