## Supplementary Info for "Convergent and selective representations of pain, appetitive processes, aversive processes, and cognitive control in the insula"

[6] Group author

* Corresponding authors

Correspondence to:

Tor D. Wager
Diana L. Taylor Distinguished Professor
Presidential Cluster in Neuroscience and
Department of Psychological and Brain Sciences
Dartmouth College


Mijin Kwon
Department of Psychological and Brain Sciences
Dartmouth College


### Supplementary Notes

#### Data harmonization and normalization

When analyzing data aggregated from multiple studies across various functional domains, it is crucial to account for two types of variability or biases to accurately identify functionally convergent and selective areas in the insula. Here we refer to these two as 1) study-wide variability and 2) functional domain-wide variability. Study-wide variability is primarily a technical issue, while domain-wide variability is both a technical and biological problem.

Study-wide variability: Data from different studies and sites are inevitably measured on different scales due to variable factors during data acquisition and analysis. These factors include scanner, acquisition, and analysis variables, such as field strength, TR, TE, acquired voxel size, flip angle, local concentration of water in tissue, choice of baseline state, stimulus timing, physiological noise removal and filtering choices, scaling of the hemodynamic response function(s) used, model regressors and contrast weights, choice to analyze percent signal change, method for converting to percent signal change, choice to resample voxels, and contrast scaling across multiple runs.

Domain-wide variability: The magnitude of brain activity elicited by each functional domain differs. For example, pain generates more pronounced (higher activation) and widespread activity across the brain, including the insula. Without proper data harmonization to account for these differences in signal levels, it would be challenging to find valid functionally-selective areas for other domains or lead to invalid results.

##### Consideration of other data harmonization and normalization methods

To address these issues, we explored a range of data harmonization and normalization methods, including ComBat[1], L2 normalization, normalization with standard deviation as a norm, and z-scoring. ComBat has been successfully applied in various neuroimaging modalities, such as DTI[2], cortical thickness[2], volumetric T1[3], functional connectivity[4,5], and task-based fMRI[5]. However, ComBat was not applicable in our case due to the nested relationship between sites/scanning parameters and task/study. L2 normalization and normalization with standard deviation as a norm did not significantly improve data harmonization compared to raw data. Therefore, we decided to analyze relative patterns after applying z-scoring normalization at the image level across voxels to account for site, scanner, and inter-study variability while still finding meaningful differences in activation patterns within the insula between the included functional domains. Although this approach may result in the loss of some informative signals, such as pain-related activity in the posterior insula, it enables the identification of local coding patterns for each domain.

Supplementary figures

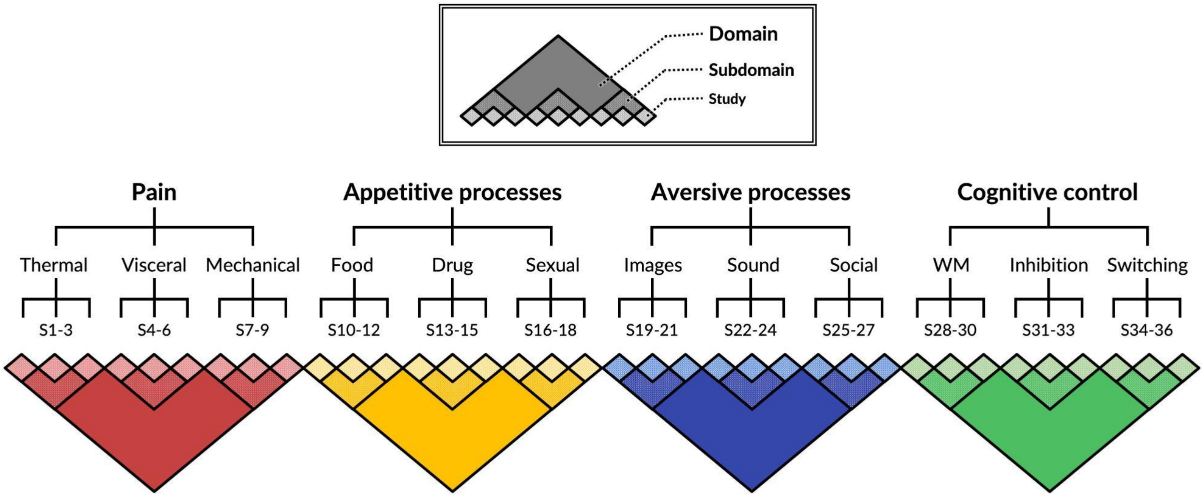

Supplementary Fig. 1

**Multi-study data structure.** Dataset used in the current study consists of 15 participants from each of 36 fMRI studies systematically sampled across four functional domains closely linked to insular function: somatic pain, non-somatic appetitive processes, non-somatic aversive processes, and cognitive control. Each functional domain comprises three subdomains representing different experimental paradigms within that domain, with three studies per subdomain to ensure representativeness and generalizability. Pain domain includes thermal (thermal stimulation), mechanical (mechanical stimulation), and visceral (visceral stimulation) subdomains. Appetitive processes domain includes food (food images), drug (drug images), and sexual (sexual images) subdomains. Aversive processes domain includes images (negative images), sound (aversive sounds), and social (negative social interactions) subdomains. Cognitive control domain includes WM (working memory), inhibition (response inhibition), and switching (attention switching) subdomains.

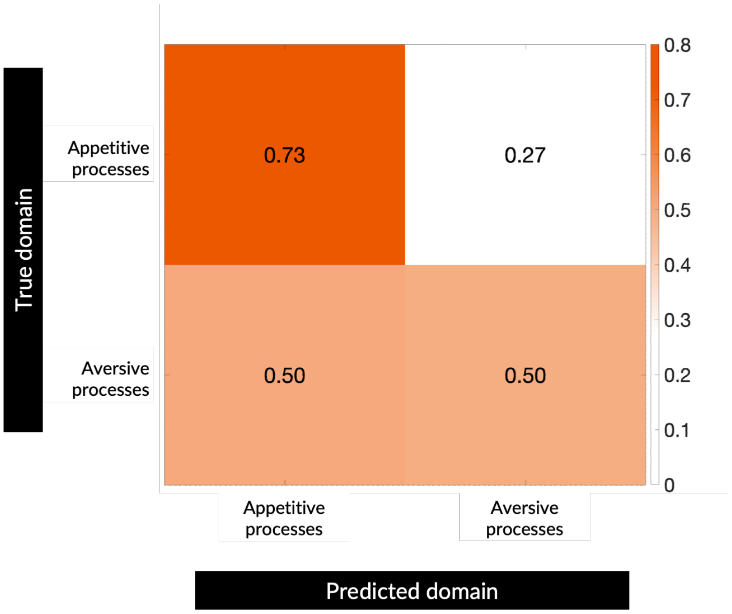

Supplementary Fig. 2

**Pairwise classification between appetitive and aversive processes.** To further investigate the multiclass classification patterns observed in Fig. 2b, we trained binary SVM classifiers specifically between appetitive and aversive processes. When classifying appetitive processes against aversive ones, accuracy was above chance (73%), but when classifying aversive against appetitive processes, accuracy was at chance level (50%). This pattern suggests that while appetitive processes have distinct neural representations, aversive processes share substantial neural patterns with a subset of appetitive processes or potentially show greater heterogeneity in their neural representations, which may result in more conservative results in identifying aversive-selective zones.

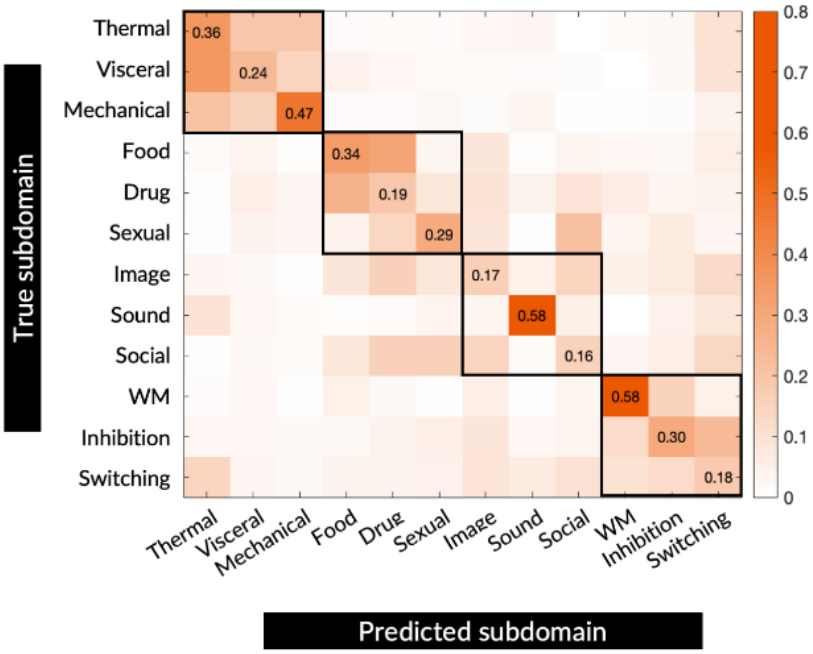

Supplementary Fig. 3

**Subdomain classification using insular activation patterns.** Confusion matrix shows prediction accuracies for multiclass SVM classifiers trained to discriminate between 12 subdomains (3 per domain) using the same leave-one-study-out scheme as the domain classification (see Methods for details). Classifier performed above chance (8.3% accuracy) for all subdomains (mean=32.11%, range: 15.6% for aversive social interaction to 57.8% for working memory), with mechanical pain, aversive sound, and working memory showing higher discriminability compared to other subdomains.

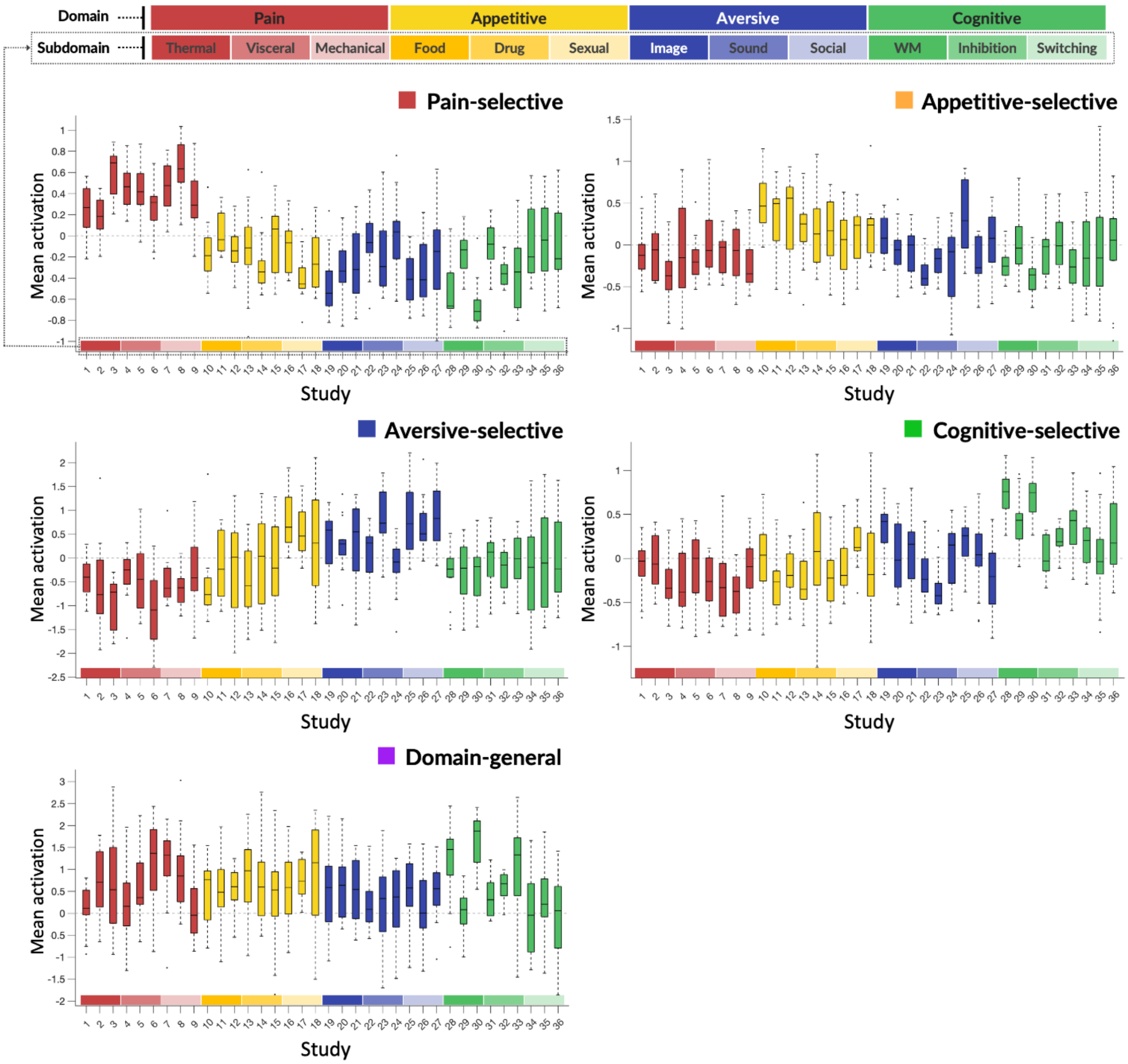

**Supplementary Fig. 4**

**Activation profiles of insular zones.** Mean z-scored contrast coefficients for domain-general and domain-selective zones across all studies, grouped by domains and subdomains. Domain-general zones show high activation across all domains, while domain-selective zones show high activation for their designated domain and low/no activation for other domains. These patterns are consistent across studies and subdomains within each domain with few exceptions. The centerline shows the median; box edges represent first (25th percentile) and third (75th percentile) quartiles, with the box length showing the interquartile range (IQR, middle 50% of data). Whiskers extend to the most extreme values within 1.5 × IQR from the box edges, and points beyond the whiskers indicate outliers.

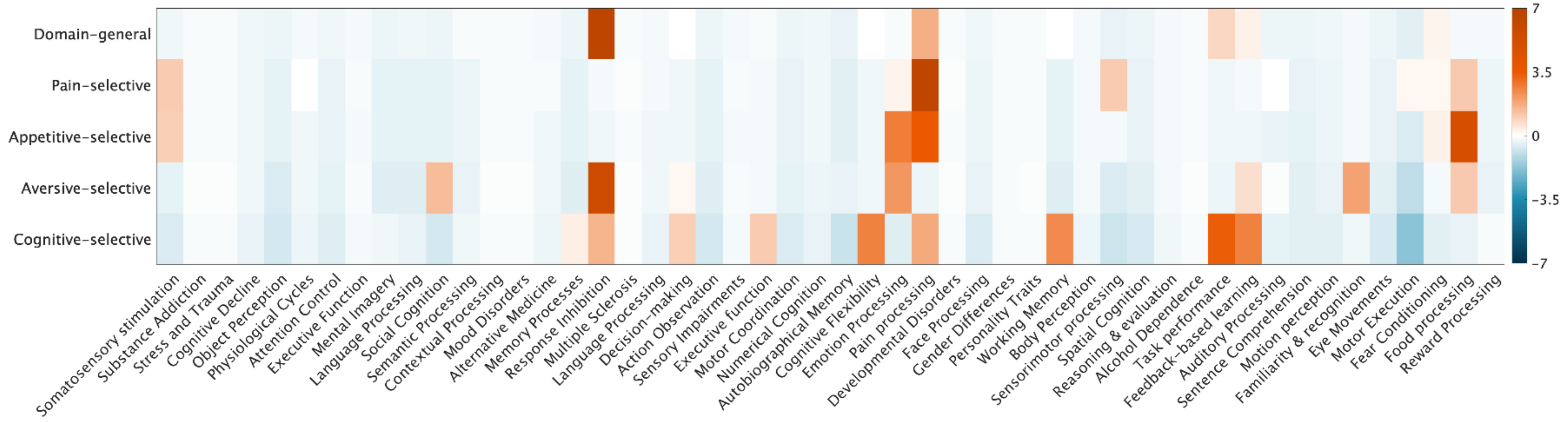

Supplementary Fig. 5

**Complete topic-level decoding of insular zones using Neurosynth.** Heatmap shows standardized point-biserial correlations between domain-general and domain-selective insular zones and all 50 psychological topic maps.

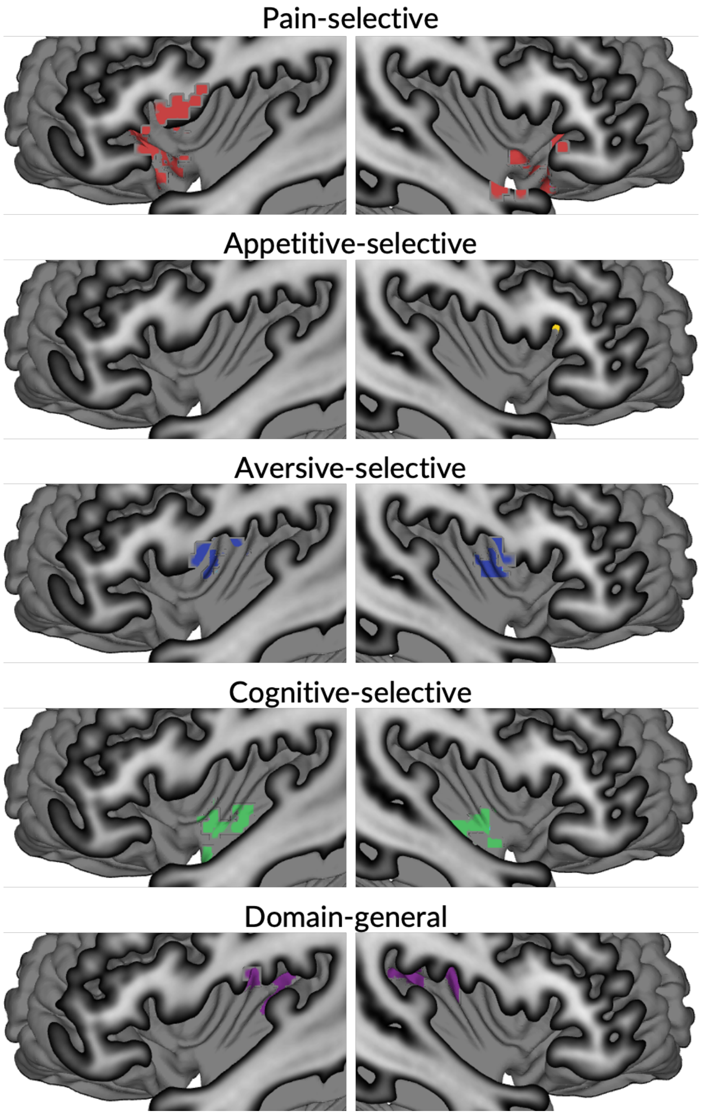

Supplementary Fig. 6

**Deactivation-based functional zones in the insula.** Domain-general zones identified from deactivation patterns were predominantly located in the posterior-most portion of the insula. Domain-selective deactivation zones showed distinct distributions: pain-selective in dorsal and ventral anterior insula, appetitive-selective in left dorsal anterior insula, aversive-selective in dorsal mid insula, and cognitive control-selective in ventral mid insula.

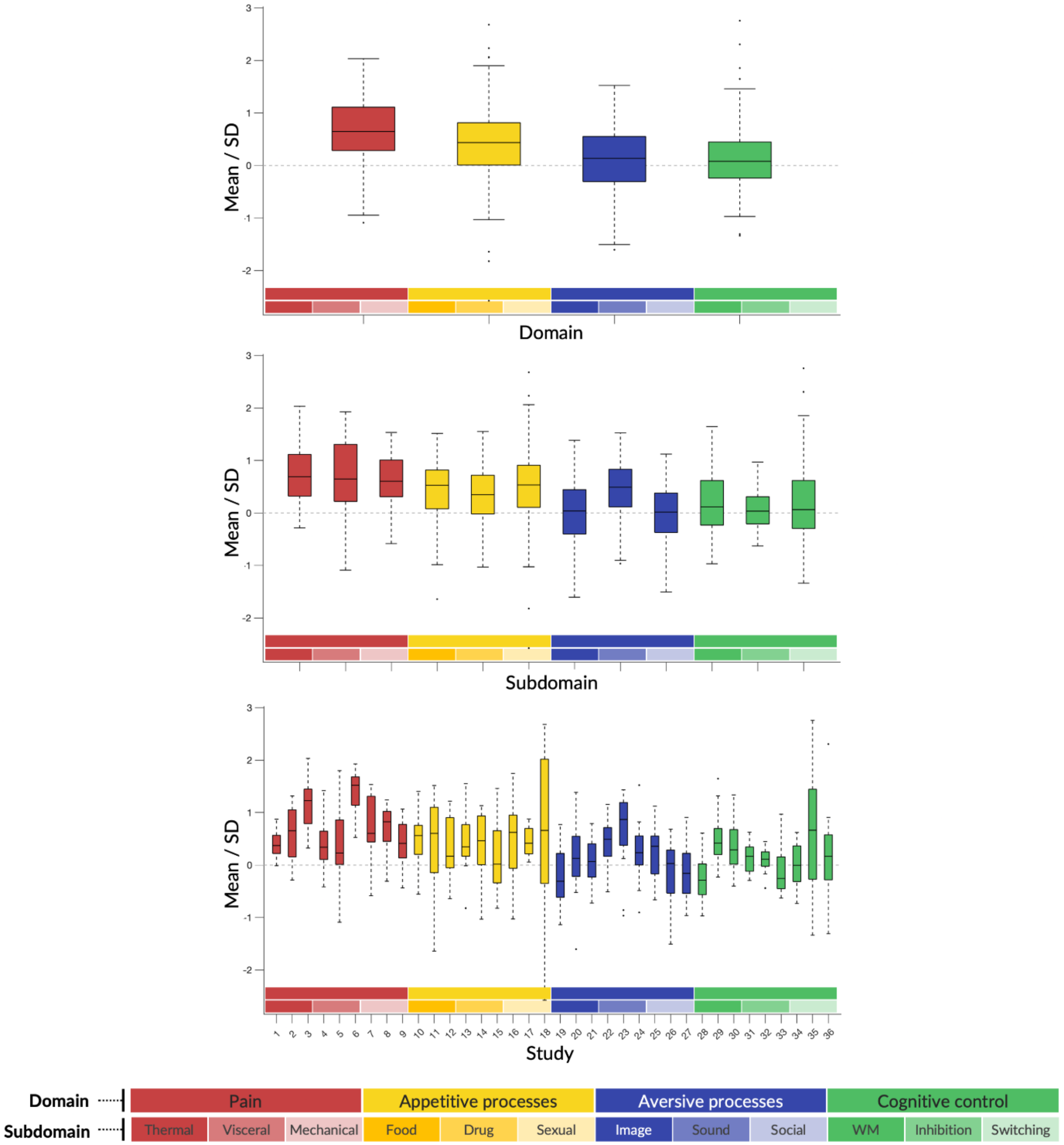

Supplementary Fig. 7

**Whole-insula activation across domains.** Standardized means (mean/SD) of contrast coefficients removed during z-score normalization shown at three levels: domains (top), subdomains within each domain (middle), and individual studies within each subdomain (bottom). All domains showed significant activation, with pain showing significantly higher standardized means compared to other domains. The centerline shows the median; box edges represent first (25th percentile) and third (75th percentile) quartiles, with the box length showing the interquartile range (IQR, middle 50% of data). Whiskers extend to the most extreme values within 1.5 × IQR from the box edges, and points beyond the whiskers indicate outliers.

1. Study selection and stratification (Repeated for all domains)

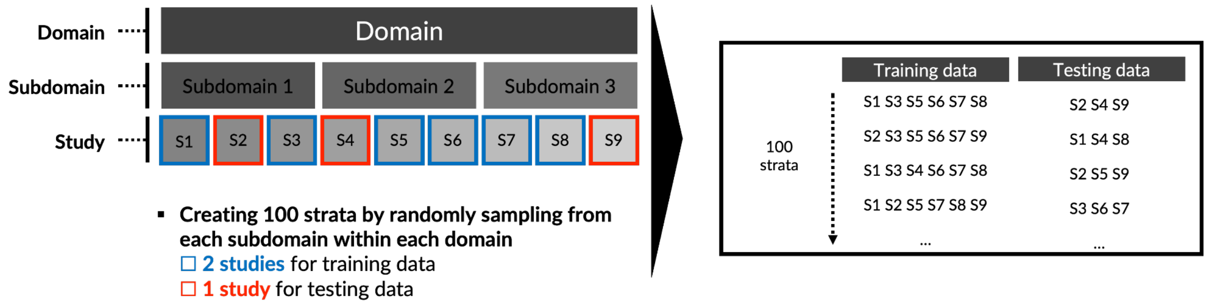

2. Model training and testing

**Training**

- Multiclass support vector machine (linear) with fitcecoc function
- One-vs-all scheme
- Hyperparameter optimization (lambda*) with 5-fold cross validation and max 5 evaluations (* *Regularization parameter inversely related to to the box constraint (C)*)

→ **Train 100 models with selected training datasets**

**Training on held-out data**

- Calculating average prediction accuracy across 100 models
- Testing significance of each domain using binomial distribution (normal approximation due to large sample size)
- Applying Bonferroni correction for multiple comparisons correction

Supplementary Fig. 8

**Schematic description of study selection and training and testing of multi-class Support Vector Machine (SVM) model.**

#### Supplementary tables

##### Supplementary Table 1

**Domain-selective and domain-general insular clusters.** Cluster information including center coordinates in standard MNI (Montreal Neurological Institute) space and parcellation labels using the following atlases: Glasser et al.[6] (HCP-MMP1.0), Failenot et al.[7] (macroanatomy), and Quab et al.[8] (cytoarchitecture).

| Insular cluster | L/R | Number of voxels | MNI Coordinates | | | Mean Bayes factors | Parcellations | | |
|---|---|---|---|---|---|---|---|---|---|
| | | | X | Y | Z | | Glasser et al. (2016) | Failenot et al. (2017) | Quab et al. (2022) |
| Pain-selective | L | 98 | -40 | 2 | 2 | 22.80 [3.08, 97.98] | Ctx_PoI2_L | Posterior short gyrus (left) | L_Area_Id6 |
| | R | 222 | 38 | -6 | 2 | | Ctx_PoI2_R | Anterior long gyrus (right) | R_Area_Op5 |
| Appetitive-selective | L | 9 | -38 | -8 | 2 | 6.88 [3.04,14.10] | Ctx_PoI2_L | Anterior long gyrus (left) | L_Area_Id5 |
| | R | 5 | 34 | -12 | -8 | | Ctx_PoI1_R | Posterior long gyrus (right) | R_Area_Ia3 |
| Aversive-selective | L | 33 | -38 | 12 | -16 | 10.12 [2.99, 25.73] | Ctx_AAIC_L | Anterior inferior gyrus (left) | L_Area_Id10 |
| | R | 42 | 36 | 14 | -14 | | Ctx_AAIC_R | Anterior inferior gyrus (right) | R_Area_TI |
| Cognitive-selective | L | 58 | -28 | 20 | 8 | 15.46 [2.96, 51.72] | Ctx_FOP4_L | Anterior short gyrus (left) | L_Area_Id7 |
| | R | 84 | 30 | 20 | 4 | | Ctx_AVI_R | Anterior short gyrus (right) | R_Area_Id6 |
| Domain-general | L | 55 | -42 | 16 | -2 | 24.60 [3.13, 68.94] (Pain)<br>26.40 [2.98, 61.20] (Appetitive)<br>11.25 [3.00, 26.22] (Aversive)<br>20.70 [3.29, 56.28] (Cognitive) | Ctx_MI_L | Anterior short gyrus (left) | L_Area_Id6 |
| | R | 65 | 44 | 18 | -4 | | Ctx_44_R | Anterior inferior gyrus (right) | R_Area_Id8' |

Supplementary Table 2

**Neurosynth topic maps selected for the current study and their correlations with each domain-general and domain-selective insular zone.**

| Topic label | Domain-selective and domain-general insular zones | | | | | Neurosynth Topic ID |
|---|---|---|---|---|---|---|
| | Pain-selective | Appetitive-selective | Aversive-selective | Cognitive-selective | Domain-general | |
| **Somatosensory stimulation** | 1.0425 | 1.0072 | -0.3993 | -0.5538 | -0.2708 | 0 |
| **Substance Addiction** | -0.1215 | -0.1551 | -0.0977 | -0.1420 | -0.1541 | 2 |
| **Stress and Trauma** | -0.1194 | -0.1531 | -0.0945 | -0.1377 | -0.1529 | 4 |
| **Cognitive Decline** | -0.2390 | -0.2606 | -0.2701 | -0.3774 | -0.2204 | 5 |
| **Object Perception** | -0.3890 | -0.3947 | -0.5712 | -0.6783 | -0.3045 | 6 |
| **Physiological Cycles** | -0.0137 | -0.2308 | -0.2214 | -0.3109 | -0.2017 | 7 |
| **Attention Control** | -0.3467 | -0.3574 | -0.4281 | -0.4966 | -0.2244 | 9 |
| **Executive Function** | -0.1460 | -0.1770 | -0.1335 | -0.1910 | -0.1679 | 13 |
| **Mental Imagery** | -0.3938 | -0.3861 | -0.4971 | -0.2657 | -0.3077 | 14 |
| **Language Processing** | -0.4150 | -0.4049 | -0.5280 | -0.3580 | -0.2688 | 15 |
| **Social Cognition** | -0.3836 | -0.3900 | 1.3726 | -0.6613 | -0.3015 | 17 |
| **Semantic Processing** | -0.2892 | -0.3057 | -0.3373 | -0.2522 | -0.1610 | 20 |
| **Contextual Processing** | -0.1155 | -0.1497 | -0.0889 | -0.1300 | -0.1508 | 21 |
| **Mood Disorders** | -0.1161 | -0.1502 | -0.0897 | -0.1311 | -0.1511 | 22 |
| **Alternative Medicine** | -0.1163 | -0.2323 | -0.2238 | -0.3142 | -0.2026 | 23 |
| **Memory Processes** | -0.4258 | -0.4058 | -0.5440 | 0.3508 | -0.3257 | 24 |
| **Response Inhibition** | -0.2176 | -0.2414 | 5.5929 | 1.5550 | 6.5496 | 25 |
| **Multiple Sclerosis** | -0.1042 | -0.1395 | -0.0723 | -0.1073 | -0.1444 | 30 |
| **Language Processing** | -0.2166 | -0.2427 | -0.2408 | -0.3374 | -0.2027 | 32 |
| **Decision-making** | -0.2214 | -0.2448 | 0.1886 | 0.9870 | -0.0404 | 34 |
| **Action Observation** | -0.3908 | -0.3970 | -0.4927 | -0.6813 | -0.3059 | 40 |
| **Sensory Impairments** | -0.1578 | -0.1876 | -0.1509 | -0.2147 | -0.1746 | 46 |
| **Executive function** | -0.1857 | -0.2149 | -0.1954 | 1.0597 | -0.1109 | 49 |
| **Motor Coordination** | -0.3480 | -0.3589 | -0.4304 | -0.5963 | -0.2820 | 51 |

| | | | | | | |
|---|---|---|---|---|---|---|
| **Numerical Cognition** | -0.2428 | -0.2640 | -0.2755 | -0.3779 | -0.2225 | 52 |
| **Autobiographical Memory** | -0.4722 | -0.4702 | -0.3787 | -0.8443 | -0.3683 | 56 |
| **Cognitive Flexibility** | -0.2249 | -0.2479 | -0.2493 | 2.6202 | 0.0488 | 58 |
| **Emotion Processing** | 0.2267 | 2.7119 | 2.1519 | -0.5264 | -0.1547 | 60 |
| **Pain processing** | 6.5060 | 3.5111 | -0.2935 | 1.7713 | 1.6717 | 61 |
| **Developmental Disorders** | -0.1082 | -0.1430 | -0.0781 | -0.1153 | -0.1466 | 62 |
| **Face Processing** | -0.3056 | -0.3192 | -0.3676 | -0.5723 | -0.2579 | 65 |
| **Gender Differences** | -0.1306 | -0.1599 | -0.1110 | -0.1602 | -0.1593 | 66 |
| **Personality Traits** | -0.1103 | -0.1450 | -0.0812 | -0.1195 | -0.1478 | 67 |
| **Working Memory** | -0.4072 | -0.4117 | -0.5167 | 2.4255 | -0.0402 | 68 |
| **Body Perception** | -0.1919 | -0.2182 | -0.2009 | -0.2829 | -0.1938 | 70 |
| **Sensorimotor processing** | 1.0622 | -0.1656 | -0.5678 | -0.7829 | -0.3348 | 72 |
| **Spatial Cognition** | -0.3599 | -0.3692 | -0.4474 | -0.6150 | -0.2885 | 75 |
| **Reasoning & evaluation** | -0.1808 | -0.2083 | -0.1846 | -0.2607 | -0.1876 | 77 |
| **Alcohol Dependence** | -0.1240 | -0.1573 | -0.1013 | -0.1470 | -0.1555 | 81 |
| **Task performance** | -0.2436 | -0.2647 | -0.2767 | 3.3536 | 0.8578 | 82 |
| **Feedback-based learning** | -0.2169 | -0.2407 | 0.7106 | 2.6108 | 0.2978 | 85 |
| **Auditory Processing** | -0.0534 | -0.3679 | -0.0616 | -0.4330 | -0.2877 | 86 |
| **Sentence Comprehension** | -0.3753 | -0.3831 | -0.4516 | -0.4684 | -0.2941 | 87 |
| **Motion perception** | -0.2930 | -0.3092 | -0.3493 | -0.4855 | -0.2508 | 88 |
| **Familiarity & recognition** | -0.1762 | -0.2041 | 1.9528 | -0.2515 | -0.1849 | 90 |
| **Eye Movements** | -0.3571 | -0.3667 | -0.4432 | -0.6138 | -0.2869 | 93 |
| **Motor Execution** | 0.1785 | -0.5495 | -1.0050 | -1.7706 | -0.4728 | 95 |
| **Fear Conditioning** | 0.1967 | 0.3050 | -0.2151 | -0.4761 | 0.2336 | 97 |
| **Food processing** | 1.1238 | 4.9280 | 1.1365 | -0.3482 | -0.2116 | 98 |
| **Reward Processing** | -0.2900 | -0.3177 | -0.3428 | -0.1450 | -0.1861 | 99 |

Supplementary Table 3

**List of highly matching cytoarchitectonic parcels for each insular zone**

| Insular cluster | L/R | Parcel | Dice coefficient | Cytoarchitectonic feature |
|---|---|---|---|---|
| Pain-selective | L | Id6 (L) | 0.260 | Dysgranular, Dorsal anterior |
| | | Id3 (L) | 0.160 | Dysgranular, Dorsal anterior |
| | | Id5 (L) | 0.120 | Agranular-dysgranular, Inferior posterior |
| | R | Id6 (R) | 0.190 | Dysgranular, Dorsal anterior |
| | | Id3 (R) | 0.160 | Granular-dysgranular, Posterior |
| | | Id2 (R) | 0.150 | Granular-dysgranular, Posterior |
| | | Ig2 (R) | 0.120 | Granular-dysgranular, Posterior |
| Appetitive-selective | L | Id5 (L) | 0.200 | Agranular-dysgranular, Inferior |
| | R | Ia3 (R) | 0.110 | Agranular, Ventral anterior cluster |
| Aversive-selective | L | Id10 (L) | 0.170 | Agranular, Ventral anterior |
| | R | Id10 (R) | 0.170 | Agranular, Ventral anterior |
| | | Id9 (R) | 0.140 | Agranular, Ventral anterior |
| Cognitive-selective | L | Id7 (L) | 0.260 | Dysgranular, Dorsal anterior |
| | | Id6 (L) | 0.120 | Dysgranular, Dorsal anterior |
| | R | Id6 (R) | 0.210 | Agranular, Ventral anterior |
| | | Id8 (R) | 0.160 | Dysgranular, Dorsal anterior |
| | | Id7 (R) | 0.150 | Dysgranular, Dorsal anterior |
| | | Id10 (R) | 0.100 | Agranular, Ventral anterior |
| Domain-general | L | Id6 (L) | 0.260 | Dysgranular, Dorsal anterior |
| | R | Id8 (R) | 0.160 | Agranular, Ventral anterior |

Supplementary Table 4

Study info

| Study # | Domain | Subdomain | Publication | N | Contrasts | Stimulus/ Paradigm | N (female) | Mean Age | IRB/Ethics Approval Committee | MRI System |
|---|---|---|---|---|---|---|---|---|---|---|
| 1 | Pain | Thermal | Atlas et al. (2010) | 15 | High vs low pain | Thermal stimulation | 19 (9) | 25.5 | Columbia University | 1.5T GE Signa TwinSpeed Excite HD |
| 2 | Pain | Thermal | Wager et al. (2013) | 15 | 49.3° C vs baseline | Thermal stimulation | 33 (22) | 27.9 | Columbia University | 1.5T GE Signa TwinSpeed Excite HD |
| 3 | Pain | Thermal | Krishnan et al. (2016) | 15 | High pain (48 degree) vs baseline | Cutaneous thermal pain | 28 (10) | 25.2 | University of Colorado Boulder | 3T Siemens Tim Trio |
| 4 | Pain | Visceral | Kano et al. (2017) | 15 | Distension vs baseline | Rectal distention | 29 (15) | 22.5 | Tohoku University School of Medicine | 3T Siemens TrioTIM |
| 5 | Pain | Visceral | Rubio et al. (2015) | 15 | Distension vs baseline | Rectal distention | 15 (9) | 24* | Comité de Protection des Personnes Sud Est V, France | 3T Philips Achieva TX |
| 6 | Pain | Visceral | Coen et al. (2011) | 15 | Esophageal distension vs rest | Esophageal pain | 31 (16) | 30 | King's College London, UK | 3T GE Signa Excite II |
| 7 | Pain | Mechanical | Unpublished | 15 | 7 kg/cm2 vs baseline | Pressure Stimulation | 15 (4) | 26.9 | University of Colorado Boulder | 3T Siemens TrioTIM |
| 8 | Pain | Mechanical | Čeko et al. (2022) | 15 | 4, 5, 6 and 7 kg/cm2 vs baseline | Pressure Stimulation | 15 (8) | 24.2 | University of Colorado Boulder | 3T Siemens Prisma |
| 9 | Pain | Mechanical | Ashar et al. (Unpublished) | 15 | High vs low pressure | Pressure Stimulation | 141 (75) | 41.7 | University of Colorado Boulder | 3T Siemens Prisma |

| | | | | | | | | | | |
|---|---|---|---|---|---|---|---|---|---|---|
| 10 | Appetitive Process | Food | Koban et al. (2023) | 15 | Food cue vs. baseline (HC*) | Craving regulation | 22 (9) | 26.4 | Columbia University | 1.5T GE Signa TwinSpeed Excite HD |
| 11 | Appetitive Processes | Food | Koban et al. (2023) | 15 | Food cue vs. baseline (HC*) | Craving regulation | 18 (6) | 42.1 | Yale University | 3T Siemens Magnetom Trio |
| 12 | Appetitive Process | Food | Koban et al. (2023) | 15 | Food cue vs. baseline (Smoker) | Craving regulation | 21 (8) | 26.8 | Columbia University | 1.5T GE Signa TwinSpeed Excite HD |
| 13 | Appetitive Process | Drug | Koban et al. (2023) | 15 | Drug cue vs. baseline (Drinker) | Craving regulation | 17 (7) | 33.4 | Yale University | 3T Siemens Tim Trio |
| 14 | Appetitive Process | Drug | Koban et al. (2023) | 15 | Drug cue vs. baseline (Cocaine user) | Craving regulation | 21 (3) | 43.5 | Yale University | 3T Siemens Magnetom Trio |
| 15 | Appetitive Process | Drug | Koban et al. (2023) | 15 | Drug cue vs. baseline (Smoker) | Craving regulation | 21 (8) | 26.8 | Columbia University | 1.5T GE Signa TwinSpeed Excite HD |
| 16 | Appetitive Process | Sexual | Wehrum et al. (2013) | 15 | Sexual vs. neutral pictures | Sexually arousing images | 100 (50) | 25.4 | German Psychologica l Society | 1.5T Siemens Symphony with quantum gradient system |
| 17 | Appetitive Process | Sexual | Stark et al. (2019) | 15 | Sexual video vs. baseline | Sexually arousing videos | 70 (33) | 25.7 | German Psychologica l Society | 3T Siemens Prisma |
| 18 | Appetitive Process | Sexual | Kragel et al. (2019) | 15 | Sexual images vs. baseline | Sexually arousing images from IAPS and GAPED | 18 (10) | 25 | University of Colorado Boulder | 3T Siemens Healthcare |
| 19 | Aversive Process | Visual | Gianaros et al. (2014) | 15 | Negative pictures vs baseline | Images from IAPS | 183 (88) | 42.7 | University of Pittsburgh | 1.5 and 3T GE Signa LX Horizon Echospeed, |
| 20 | Aversive Process | Visual | Yarkoni et al. (2011) | 15 | Negative vs neutral pictures | Images from IAPS | 108 (NR) | NR | Stanford University, Columbia University | 1.5T GE Signa Twin Speed Excite HD scanner |

| | | | | | | | | | | |
|---|---|---|---|---|---|---|---|---|---|---|
| 21 | Aversive Process | Visual | Kober et al. (2019) | 15 | Negative pictures vs baseline | Images from IAPS | 16 (5) | 31.75 | Columbia University | 1.5T GE Signa Twin Speed Excite HD scanner |
| 22 | Aversive Process | Auditory | Čeko, Woo et al. (Unpublished) | 15 | Unpleasant Sounds vs baseline | Sounds from IADS | 15 (7) | 31.1 | University of Colorado Boulder | 3T Siemens Tim Trio |
| 23 | Aversive Process | Auditory | Čeko et al. (2022) | 15 | Unpleasant Sounds vs baseline | Sounds from IADS | 15 (9) | 24.4 | University of Colorado Boulder | 3T Siemens Tim Trio |
| 24 | Aversive Process | Auditory | Ashar et al. (Unpublished) | 15 | Sound high vs. low in unpleasantness | Aversive sound (knife scraping on glass) | 141 (75) | 41.7 | University of Colorado Boulder | 3T Siemens Prisma |
| 25 | Aversive Process | Social | Kross et al. (2011) | 15 | Images of ex-partner vs friend | Images of ex-partners | 40 (21) | 20.8 | Columbia University | 1.5T GE Signa TwinSpeed |
| 26 | Aversive Process | Social | Krishnan et al. (2016) | 15 | High pain pictures vs baseline | Images of others in pain | 30 (12) | 25.2 | University of Colorado Boulder | 3T Siemens Tim Trio |
| 27 | Aversive Process | Social | Yu et al. (2020) | 15 | Self-incorrect vs. baseline | Guilt from causing pain due to one's error | 24 (11) | 22 | Peking University | 3T Siemens Tesla Trio |
| 28 | Cognitive Control | WM | DeYoung et al. (2009) | 15 | 3-back blocks vs baseline | N-back (faces and words) | 104 (59) | 22.7 | Washington University Medical Center | 3T Siemens Allegra |
| 29 | Cognitive Control | WM | van Ast et al. (2016) | 15 | N-back blocks vs baseline | N-back (words) | 21 (10) | 22.2 | Columbia University | 3T Philips Achieva |
| 30 | Cognitive Control | WM | Unpublished | 15 | Word event vs. fixation | Updating working memory task | 30 (16) | 28.1 | University of Colorado Boulder | 3T Magnetom Trio |
| 31 | Cognitive Control | Inhibition | Aron et al. (2007) | 15 | All trials vs baseline | Stop signal Task | 15 (5) | 28.1 | UCLA | 3T Siemens Allegra |
| 32 | Cognitive Control | Inhibition | Xue et al. (2008) | 15 | All trials vs baseline | Stop signal Task | 15 (9) | 23.6 | UCLA | 3T Siemens Allegra |
| 33 | Cognitive Control | Inhibition | Unpublished | 15 | Antisaccade vs. fixation | Response inhibition task | 30 (16) | 28.1 | University of Colorado Boulder | 3T MAGNETOM Trio |
| 34 | Cognitive Control | Switching | Unpublished | 15 | Switching vs. fixation | Set shifting task | 30 (16) | 28.1 | University of Colorado Boulder | 3T MAGNETOM Trio |

| | | | | | | | | | | |
|---|---|---|---|---|---|---|---|---|---|---|
| 35 | Cognitive Control | Switching | Wager et al. (2005) | 15 | Switch vs. non-switch | Attention switching task | 39 (NR) | NR | University of Michigan | 3T GE Signa |
| 36 | Cognitive Control | Switching | Wager et al. (2005) | 15 | Switch vs. non-switch | Attention switching task | 39 (NR) | NR | University of Michigan | 3T GE Signa |

#### Reference

1. Johnson WE, Li C, Rabinovic A. Adjusting batch effects in microarray expression data using empirical Bayes methods. *Biostatistics*. 2007;8(1):118-127. doi:10.1093/biostatistics/kxj037
2. Fortin JP, Parker D, Tunç B, et al. Harmonization of multi-site diffusion tensor imaging data. *NeuroImage*. 2017;161:149-170. doi:10.1016/j.neuroimage.2017.08.047
3. Pomponio R, Erus G, Habes M, et al. Harmonization of large MRI datasets for the analysis of brain imaging patterns throughout the lifespan. *NeuroImage*. 2020;208:116450. doi:10.1016/j.neuroimage.2019.116450
4. Yu M, Linn KA, Cook PA, et al. Statistical harmonization corrects site effects in functional connectivity measurements from multi-site fMRI data. *Hum Brain Mapp*. 2018;39(11):4213-4227. doi:10.1002/hbm.24241
5. Nielson DM, Pereira F, Zheng CY, et al. Detecting and harmonizing scanner differences in the ABCD study - annual release 1.0. Published online May 2, 2018:309260. doi:10.1101/309260
6. Glasser MF, Coalson TS, Robinson EC, et al. A multi-modal parcellation of human cerebral cortex. *Nature*. 2016;536(7615):171-178. doi:10.1038/nature18933
7. Faillenot I, Heckemann RA, Frot M, Hammers A. Macroanatomy and 3D probabilistic atlas of the human insula. *NeuroImage*. 2017;150:88-98. doi:10.1016/j.neuroimage.2017.01.073
8. Quabs J, Caspers S, Schöne C, et al. Cytoarchitecture, probability maps and segregation of the human insula. *NeuroImage*. 2022;260:119453. doi:10.1016/j.neuroimage.2022.119453
